# Climatic conditions predict embryonic development in thorn tailed Rayadito (*Aphrastura spinicauda*)

**DOI:** 10.64898/2026.03.11.710834

**Authors:** Joseph Badji-Churchill, Martje Birker-Wegter, Maaike Versteegh, Rodrigo Vàsquez, Jan Komdeur

## Abstract

Rapid changes to weather caused by climate change have a negative effect on much of the worlds animal populations and species. Some populations are more vulnerable than others to the effects of climate change, and individuals are particularly vulnerable during early development. Good embryonic development is important for vertebrate species because this can dictate their breeding success and survival rates, and disruptions to this phase can have far reaching fitness effects that can last into adulthood and beyond. We looked at the impact of weather (ambient temperature, rainfall and wind speed) on the embryonic development of thorn-tailed Rayadito (*Aphrastura spinicauda*) at two different latitudes in Patagonia, Chile. We measured the heart rate of embryos just before hatching using an egg buddy machine to determine embryonic development. Optimum development of nestlings is important for fledging, so it is essential that embryonic development is successful. We studied two populations. One is situated in a temperate rainforest on the northern border of Patagonia called Pucon which we studied in 2018 and 2019, with mild temperatures (12 degrees Celsius), high rainfall (636ml) and low wind speeds (6.3km/h). The other is in a sub-Antarctic old growth forest in southern Patagonia called Navarino island which we studied in 2018, 2019 and 2023, which is comparatively drier (138ml), colder (8.3 degrees Celsius) and has higher average wind speeds (16.6km/h). We found that embryonic development was better in the south compared to the north, indicated by higher embryo heart rates near hatching in the south. Development of embryos in the northern site was slower when conditions were cold and windy. Development of embryos in the southern site was unaffected by temperature, rainfall or wind speeds. In northern Patagonia, when minimum temperatures were low and wind speeds high during the period encompassing clutch completion, initiation of full incubation and during incubation, have a negative impact on embryonic development. In contrast, when Rayaditos in the southern population experience slow embryonic development, they extend the incubation period allowing embryos more time to develop before hatching. Our study shows that in the north of Patagonia embryonic development declines over years and that Rayaditos seem not to have adapted to dealing with climate change. On the other hand, in the south of Patagonia, embryonic development is unaffected by climatic factors, suggesting that Rayaditos are adapting to climate change through extending their incubation periods, allowing embryos to fully develop before hatching. It appears that Rayaditos in the northern population are not extending their incubation periods and are not adapting to the threats posed by climate change. To our knowledge, this is the first study of its kind to examine embryonic development in the field and to associate this to changing weather patterns whilst highlighting specific days on which development is influenced.

## INTRODUCTION

The planet’s animal populations are struggling to adapt to the conditions caused by breakdowns in climate (Habibullah et al., 2022; Kabir et al., 2023; Wudu et al., 2023). Changes to our planet’s weather presents new selection pressures for populations and poses serious risk to animal population dynamics. Climatic selection pressures are most impactful during key developmental phases, like embryonic development (Sirard, 2021; Jonsson, Jonsson & Hansen, 2022; Tona et al., 2022). Birds are particularly vulnerable to climate change during embryonic development compared to most mammals, because they develop in eggs outside of the mother’s womb. The condition and development of avian embryos are influenced by weather conditions, but other factors like incubation patterns, nest quality, clutch size, parental quality and genetics are also important (Yahav and Brake, 2014; Du et al., 2019; Nord and Giroud, 2020). Embryonic development can be measured through several indices. Measuring embryonic heart rate is the simplest, least invasive and most utilisable in the field. Other indices for embryonic development, such as egg mass, yolk components and hormone levels have also been used, but these are often expensive and invasive techniques and egg mass fluctuates too much to be reliable in isolation (Mentesana et al., 2019: Du et al., 2019; Nord and Giroud, 2020). Embryonic heart rate can be reliably measured throughout the entire development using an egg buddy machine (Vetronic Services, Abbotskerswell, Devon, UK).

The effect of temperature on embryonic development in lab-controlled conditions is well documented. It is commonly found that lower temperatures correlate with lighter hatchlings, longer developmental periods and poorer thermoregulation (Olson et al., 2006; Nord and Giroud, 2020). These lab studies often keep the incubation temperature constant throughout observations, which is not the case in the wild where temperatures fluctuate. Therefore, it is important to investigate if there are some stages of embryonic development where embryos are more vulnerable than others, or if temperature changes at certain times effect embryos more than others. Field studies are rare and do not address specific time periods during which climatic factors can influence embryonic development (Du and Shine 2022). One field study that captures the impact of temperature on embryonic development succinctly is by Griffith et al., (2016) on Australian zebra finches (*Taeniopygia guttata*), where increases of 6 Celsius degrees in nest temperatures caused eggs to hatch 3% quicker and so accelerated embryonic development. This is supported by another field study in Australia on zebra finches that found higher egg temperatures increase embryonic heart rates, which is indicative of faster embryonic development (Sheldon et al., 2018). These studies demonstrate that higher nest temperatures lead to faster embryonic development, which in turn leads to faster hatching. We also see the reverse in lab-based experiments on domestic chicken (*Gallus gallus*), zebra finch and wood duck (*Aix sponsa*) embryos, where egg cooling lengthens the incubation period and slows development, which lead to smaller nestlings or nestlings that could not thermoregulate effectively (Suarez et al., 1996, Olson, Vleck and Vleck, 2006; Durant et al., 2013). These consequences could be disadvantageous or even fatal to wild populations living in harsh conditions where individuals need the best start possible to survive their first year (Nord and Nilsson, 2016). What is important to highlight here is that these studies all focus on internal nest temperatures, whilst we focus on ambient temperatures, which is part of the novelty of our study. To gain a better understanding of the effects of weather patterns on avian embryonic development, we need to investigate a broad range of habitats differing in climatic conditions, as the effects of weather patterns on embryonic development is largely undocumented. The aim of our study is to investigate the effect of climatic factors on embryonic development, and whether parents use mitigating behaviours to prevent poor embryonic development, using the thorn-tailed Rayadito (*Aphrastura spinicauda*) as a model species.

The thorn tailed Rayadito is a non-migratory suboscine species found across Chile and Argentina (Figure 1). We use the Rayadito as a model species because it is common across a large geographical range which varies in climatic conditions, which in turn may act as selection pressures for the species to adapt to a range of environmental conditions. As such, the Rayadito is a useful indicator species for other species facing similar climatic challenges. We carried out data collection at two locations in Chile; Pucón (northern site); and Navarino (southern site) to compare the same species existing in two contrasting environments to determine how Rayaditos respond to different climate-based selection pressures. We expect embryos in our sites to respond similarly to temperature as seen elsewhere where increases or decreases in temperatures correlate with a speeding or slowing of development. Previous studies investigate climatic variables in isolation, but we consider multiple climatic factors simultaneously (ambient average temperature, minimum temperatures, maximum temperatures, rainfall and wind speeds) which we expect will complicate findings as climatic variables do not correlate at our study sites.

**Figure 1.**
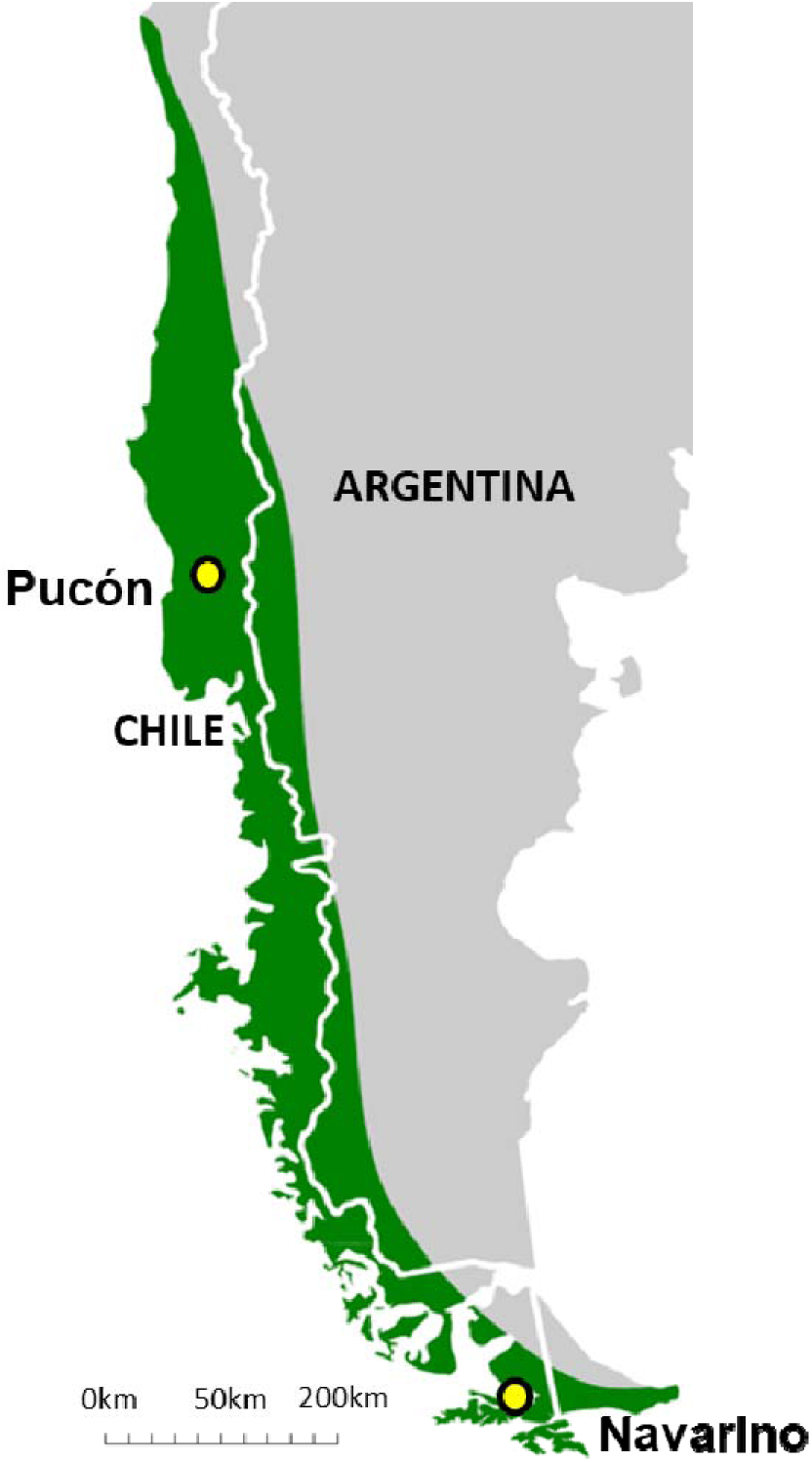
Locations of our two study sites, Navarino (southern) and Pucón (northern). The green coverage shows the distribution of the thorn-tailed Rayadito.

We recorded embryonic heart rate, as a proxy for embryonic development over three years at two different latitudes and then observed how weather impacted the heart rate close to hatching. Embryonic heart rate has been shown to be a good proxy of embryonic development, particularly in the field (*see* Du et al., 2010; Sheldon et al., 2018). Our study has two aims. First, to investigate whether climatic factors affect embryonic development in the thorn-tailed Rayadito and during what time periods is weather most influential on development. Specifically, we expect that the weather during crucial stages before embryonic development predicts embryonic heart rate. For example, the weather during the laying phase, and/or at the initiation of full incubation may explain developmental trajectories. In warmer, drier and calmer conditions we expect embryonic heart rates to be higher and in colder, wetter and windier conditions we expect heart rates to be lower. Second, if climatic factors have a negative effect on embryonic development, are parents able to use mitigating behaviours like extending incubation periods to prevent potentially poor embryonic development.

## METHODS

We studied Rayadito populations in two different latitudes in Chile that breed in nest boxes (figure 1). We collected data on Rayadito embryos during the breeding season (September to December) at two locations, Navarino island (54.932° S, 67.605° W) in 2018, 2019 and 2023 and Pucón, (39.272° S, 71.977° W) ca. 1800 km to the north in 2018 and 2019. The southern Navarino site was ca. 5km in radius, had around 200 nest boxes with 14.3% occupancy rate and were placed in 2002-3, in sub-Antarctic deciduous forest. The ambient average temperature over the breeding seasons in the years we collected data was 8.4°C (7.7°C - 9.2°C), total yearly rainfall averaged 138ml (87ml - 202ml) and wind speeds averaged 16.62km/h (15.95km/h - 17.80km/h) (INIA, 2023). Nest boxes in Navarino were dominated by deciduous species such as *Nothofagus antarctica, N. betuloides* and *N. pumilio,* whilst the understory was dominated by *Berberis mycrophylla* (evergreen shrubs). The Pucón study site had 240 nest boxes with 10.2% occupancy rate, spread out over ca. 10km radius and were placed in 2014. Pucón is dominated by a temperate rainforest with high precipitation and relatively high ambient temperatures compared to Navarino. During the breeding season the temperature averaged 12.6°C (12.5°C - 12.6°C), there was an average total yearly rainfall of 636ml (512ml - 760ml) and the wind speeds averaged 6.3km/h (6.2km/h - 6.4km/h) (INIA, 2023). In Pucón, tree species were predominantly conifer-broadleaf species such as *Lophozonia obliqua, Nothofagus dombeyi, Laurelia sempervirens, Saxegothaea conspicua* and *Laureliopsis philippiana*. The understory composition was dominated by *Chusquea spp* (Bamboo) *Rhaphithamnus spinosus* (evergreen shrub) *Azara spp* (flowering plants).

### Egg morphometrics and embryo physiology

We sampled 226 eggs (Navarino = 146, Pucón = 80) from 94 nests (Navarino = 50, Pucón = 44) in total. In Navarino, we recorded clutches of 3 (n = 4), 4 (n = 26), 5 (n = 57), 6 (n = 51) and 7 (n = 3) and in Pucón we recorded clutches of 1 (n = 5), 2 (n = 33), 3 (n = 36) and 4 (n = 6). We measured the weight (g) of each egg to the nearest 0.01g and the width and length (mm) to the nearest 0.1mm and marked each egg sequentially with dots (1, 2, 3 etc.) using non-toxic, water-based indelible ink. Rayaditos lay one egg every two days, so we visited the next day after the first egg is laid to find the second egg, or the day after that, to establish the laying pattern. Once this was established, we visited the nest every two days to measure each new egg. When no new eggs were laid and the clutch was completed (recorded as day 1), we visited the next day (three days after clutch completion) to measure the embryonic heart rate of every egg using an egg buddy heart monitor machine following standardised procedures (Vetronic Services, Abbotskerswell, Devon, UK). We measured the eggs in random order as embryonic heart rate is directly linked to temperature and egg cooling whilst eggs are being measured may impact the results (Nord and Giroud, 2020). We measured each egg from three days after clutch completion (CC+3) onwards until we detected the first heart rate and then we measured the heart rate at 12 days after clutch completion (CC+12, known from here as HR12). This is an important heart rate measurement as this is very close to hatching, with most clutches hatching 14-18 days after clutch completion. We checked the nest every day after HR12 for hatching. We recorded the number of days between clutch completion and hatching as the incubation length. In 2018 and 2019 only eggs 1 and 4 were measured for embryonic heart rate in parallel with a separate experimental study (Birker-Wegter, 2024) whilst in Navarino in 2023, all eggs were measured. We found that there are no differences in HR12 between egg numbers in Pucón (est = −1.320, std. error = 3.431, P = 0.702) or in Navarino (est = 2.644, std error = 2.920, P = 0.367), therefore we used all available data.

### Weather measurements

Temperature (°C), precipitation (accumulated ml per day) and wind speed (km/h) data were taken from meteorological stations on both sites (Table 1). In Navarino the meteorological station was a maximum of 5 km away from the nest boxes. In Pucón the station was located a maximum of 7 km from the nest boxes. The data included the daily average temperature (°C), daily minimum temperature (°C), daily maximum temperature (°C), total daily precipitation (ml) and daily average wind speed (km/h) from September 1^st^ each year – the date of the last measurement (which varied year-to-year). The daily ambient temperature and wind speed was recorded every hour for 12 hours (6 am - 6 pm) which we used to calculate the daily average. We used the accumulated precipitation for the full 24 hours.

**Table 1.**
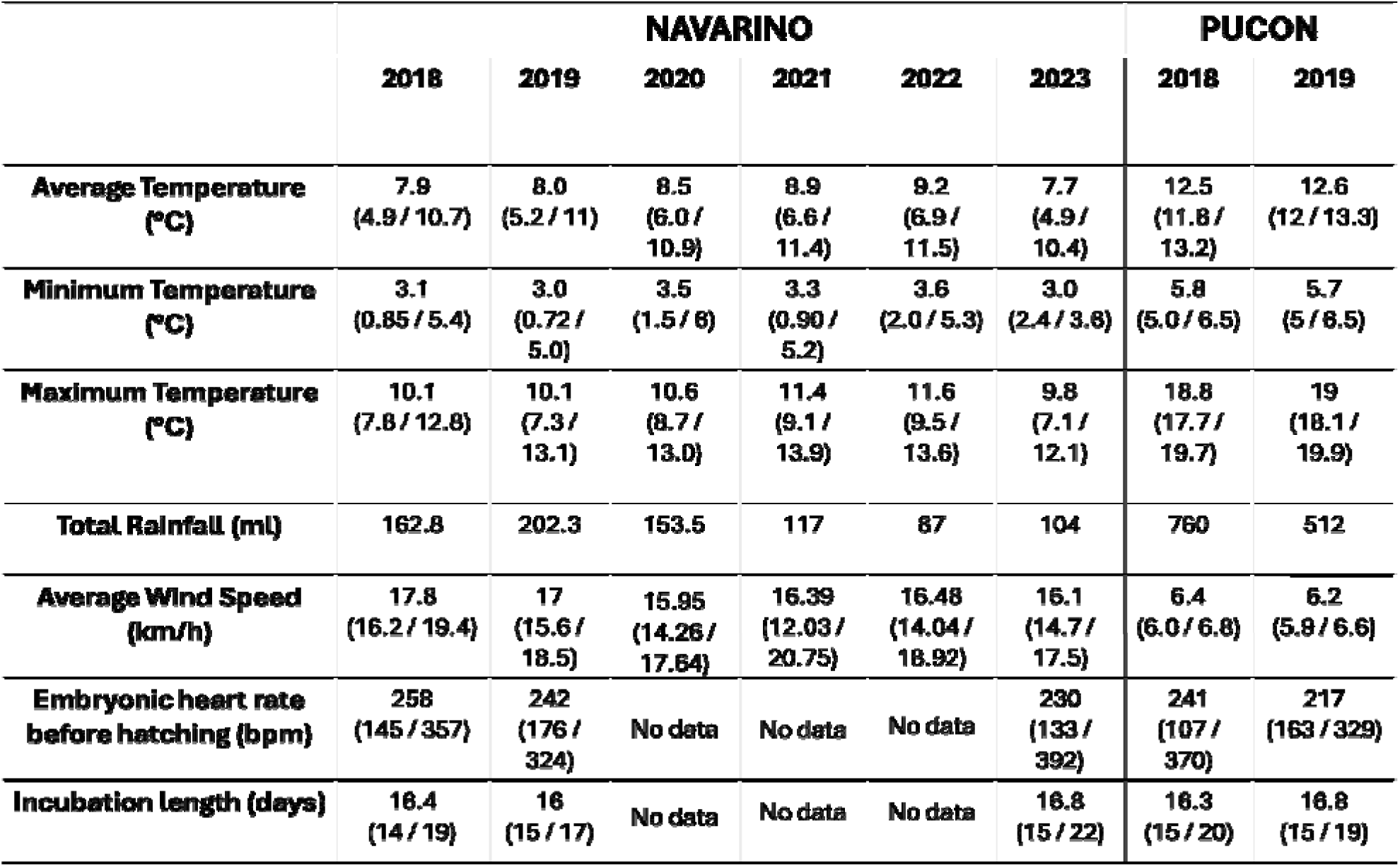
Climate, embryonic heart rates, and incubation length averages over the breeding season in each year in each location.

### Statistical analyses

HR12 was log10 transformed to achieve normal distribution (from now on referred to as logHR12). We first performed a set of t-tests to investigate differences in logHR12 and incubation length between the two locations. Because we expect populations to have developed different reaction norms to climatic factors, the rest of the analyses were done for both locations separately.

To analyse which specific days the daily ambient average, daily minimum and maximum temperature, daily total rainfall and daily average wind speed best predict embryonic heart rate, we used the ‘climwin’ package. This package correlates the effect of different climate windows on a response variable and evaluates them based on the Akaike Information Criterion (AICc) (Bailey and Van de Pol, 2016). We used embryonic HR12 as our response variable, with the weather variables as our explanatory variables and year as covariate in our baseline model (*see* Bailey and Van de Pol, 2016). All statistics were carried out using R in Rstudio v3.5.3 statistical software (Posit team, 2026). The ‘SlidingWin’ function in the climwin package was used to identify the time-window where weather factors explained logHR12 best (*for more detail see* Bailey and Van de Pol, 2016). To assess the validity and significance of the best given window, we ran 1000 random models with the same data using the ‘RandWin’ function in the climwin package. If a reliably significant result was found (P <0.05), this data was then extracted for further modelling with other predictor variables, hence not all climatic variables were explored in further modelling.

After this we ran LMER models on the logHR12 data. In Navarino, climwin identified none of the climatic to be important for logHR12. In Pucón the following climatic predictors were identified as being important; daily minimum temperature (°C), total daily rainfall (ml) and daily average wind speeds (km/h). In both Navarino as Pucón we used egg mass (g), clutch size, incubation length (days) and, in Pucón only, clutch attempt (first and second attempt) as explanatory variables. We used year as covariate and nest box number as a random factor. We also ran these models using the year and nest box number combined as a random factor to mitigate for different Rayadito pairs using the same nest boxes between years and found similar results and we do not report these results. The incubation length was measured in days from the initiation of full incubation up to hatching, which ranged 13-22 days. Because in some cases Rayadito pairs in Pucón produce more than one clutch over the breeding season and it has been observed in other species that second clutches occasionally produce smaller nestlings or nestlings with lower survival rates (Verhulst, 1998; Nager, Monaghan and Houston, 2000), the clutch attempt was recorded. We looked at the VIF scores of each model to determine that predictor variables were not linearly codependent, and residuals and dispersions were checked using residual simulation graphs and dispersion tests and conclude that data were normally distributed.

## RESULTS

### Embryonic development, incubation length and climatic variables in Pucón and Navarino

We measured embryonic development through measuring HR12 and incubation lengths (Table 1). Both in Navarino as in Pucón, all the eggs with a heart rate on day 12 hatched. We found that HR12 was significantly higher in Navarino compared to Pucón (t = 2.20, df = 105.96, P = 0.03) but incubation lengths were not significantly different between the two locations with Navarino averaging 16.6 incubation days and Pucón averaging 16.4 incubation days (t = −1.34, df = 176.18, P = 0.19).

### Climate based effects on embryonic development in Pucón and Navarino

We did not find climate-based effects on embryonic development in the population in Navarino, however we did find them near Pucón (Table 2A and 2B). In the climwin analysis, we found that minimum daily temperatures, total daily rainfall and daily wind speeds^2^ in Pucón predict embryonic development, proxied by heart rates 12 days after clutch completion (HR12) and that daily average temperature and daily maximum temperatures had no association to HR12 in Pucón. Specifically, we found that the minimum temperatures during two windows, 19-20 days and 6-13 days before HR12 were measured, were able to predict HR12. 19-20 days before HR12 correspond with the egg gestation period and 6-13 days before HR12 correspond with the period between egg laying and clutch completion, which also includes partial incubation. The relationship between minimum temperatures and HR12 is linear, with a higher minimum temperature corresponding with higher HR12 (Figure 2). We found that rainfall 1-5 days before HR12 best predicted HR12 (Figure 3). This time window is during full incubation and the day before the HR12 measurement is taken. The relationship was a negative linear correlation, where higher rainfall was associated with lower HR12. Finally, we found that wind speeds 12-13 days before HR12 best predicted HR12. This places the time window at the day of clutch completion and the initiation of full incubation, concurrent with our predictions. This showed a quadratic relationship (wind speed^2^), meaning there is an optimum wind speed for HR12. The lowest wind speeds recorded were close to 0km/h with the bulk of data points experiencing wind speeds below 2.5km/h whilst the highest wind speeds during this window were up to 12km/h and those with the highest heart rates being in the middle around 4km/h (Figure 2). In Navarino, we found that none of the climatic variables (daily average temperature, daily minimum temperature, daily maximum temperature, daily total rainfall or daily average wind speeds) correlated with HR12. Therefore, further analyses in Navarino with respect to HR12 only consisted of non-climatic factors (year, clutch size, egg mass, incubation length and nest box number).

**Figure 2-.**
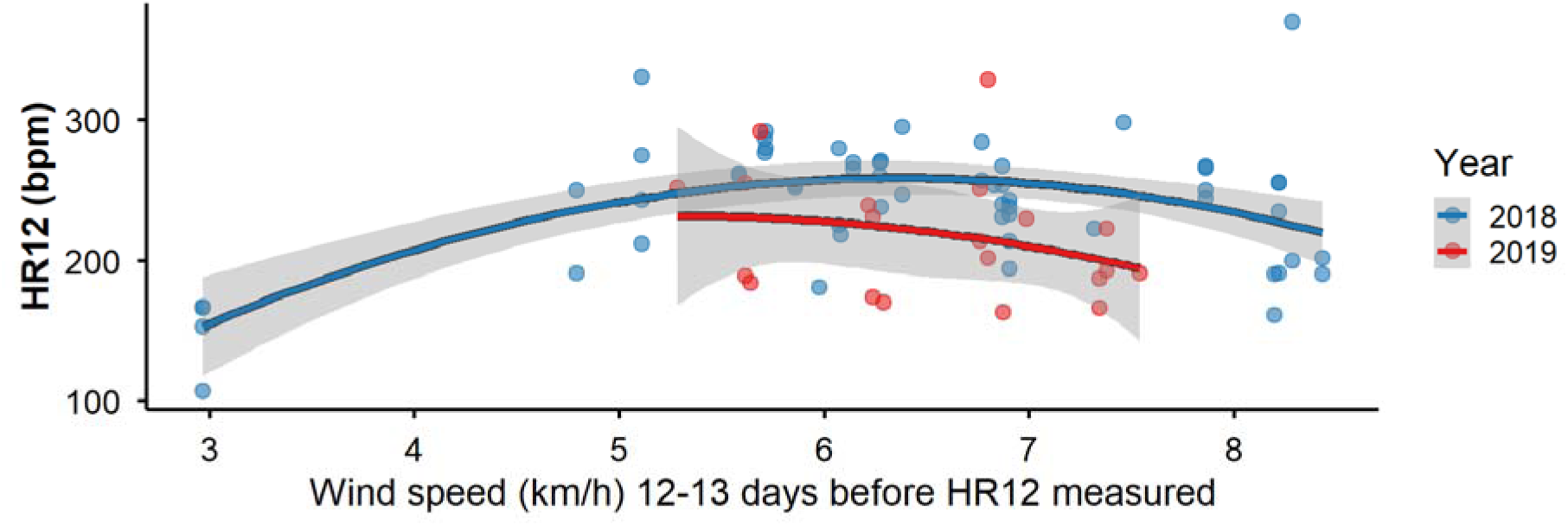
The correlation between average wind speed^2^ (km/h) 12-13 days before HR12 (heart rate at 12 of full-time incubation) in thorn tailed Rayaditos (N = 78, 2018 = 58, 2019 = 20) were measured in Pucón in 2018 & 2019. For statistical analyses, see table 2B.

**Figure 3-.**
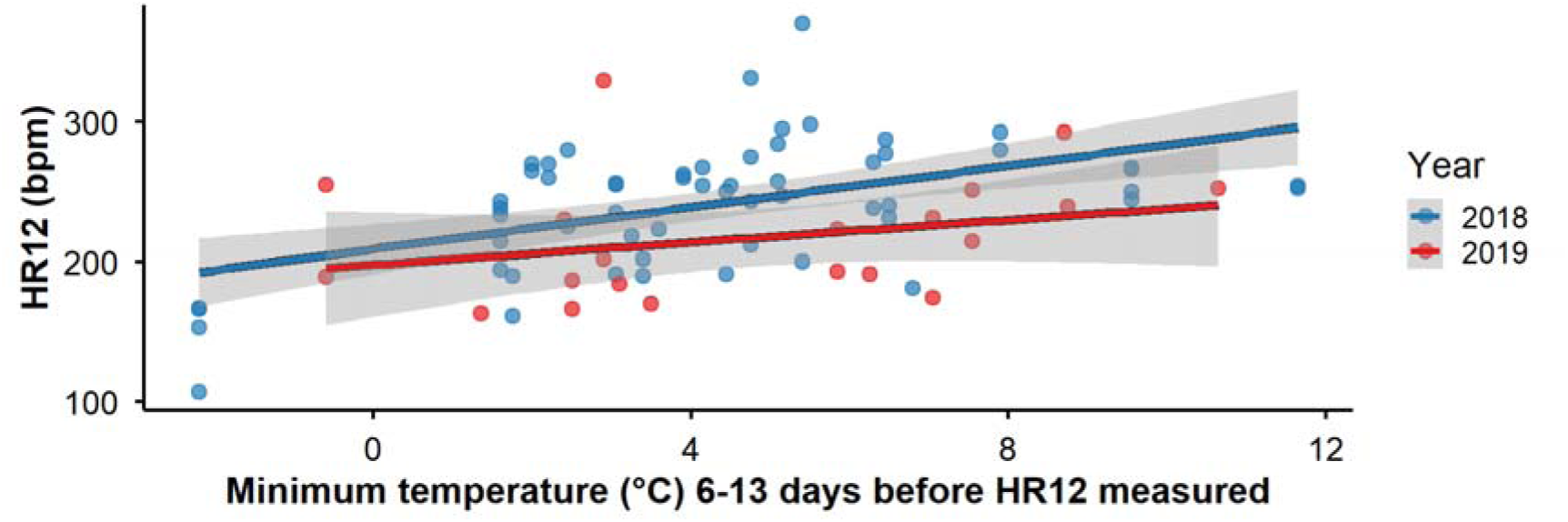
The correlation of daily minimum temperatures (°C) 6-13 days before HR12 (heart rate at 12 of full-time incubation) with HR12 in thorn tailed Rayadito (N = 78, 2018 = 58, 2019 = 20) in Pucón 2018 & 2019. For statistical analyses, see table 2B.

**Table 2-.** results of LMER models for (A) Navarino (N = 134, 2018 = 27, 2019 = 18, 2023 = 89), and (B) for Pucón (N = 78, 2018 = 58, 2019 = 20).

| Fixed effect | Estimate | SE | P-value |
| --- | --- | --- | --- |
| **YEAR (2023)** | -0.037 | 0.025 | 0.26 |
| **CLUTCH SIZE** | 0.008 | 0.012 | 0.47 |
| **EGG MASS**<br>**(g)** | 0.080 | 0.060 | 0.17 |
| **INCUBATION LENGTH**<br>**(days)** | -0.015 | 0.008 | **0.05** |

B)
| Fixed effect | Estimate | SE | P-value |
| --- | --- | --- | --- |
| **MINIMUM TEMPERATURE**<br>**(°C)** | 0.008 | 0.003 | **<0.01** |
| **ACCUMULATED RAINFALL**<br>**(ml)** | -0.0002 | 0.001 | 0.86 |
| **WIND SPEED**<br>**(km/h)** | -0.003 | 0.007 | 0.60 |
| **WIND SPEED<sup>2</sup>**<br>**(km/h)** | -0.016 | 0.005 | **<0.01** |
| **YEAR** | -0.067 | 0.020 | **<0.01** |
| **EGG MASS**<br>**(g)** | -0.025 | 0.050 | 0.51 |
| **CLUTCH SIZE** | 0.017 | 0.013 | 0.18 |
| **INCUBATION LENGTH**<br>**(days)** | -0.008 | 0.008 | 0.27 |
| **CLUTCH ATTEMPT** | -0.008 | 0.029 | 0.76 |

### Incubation length and embryonic development

In Pucón LMERs, minimum daily temperatures, daily average wind speed^2^ and year all significantly correlated with HR12 (Table 2B). Daily minimum temperature showed a strong positive linear relationship with HR12, where higher minimum temperatures predicted higher HR12. Wind speed^2^ shows a strong quadratic relationship to HR12 where moderate wind speeds returned the highest HR12 and very low or very high wind speeds resulted in lower HR12. Rainfall is not significant in predicting HR12 upon further analysis in LMERs. Additionally, we found that rainfall, egg mass, clutch size, incubation length and clutch attempts had no impact on HR12. In Navarino LMERs, we found that incubation length was the best predictor of HR12 where eggs with longer incubation lengths had lower HR12 (Figure 4 and Table 2A). Which means that year, clutch size and egg mass had no association to HR12.

**Figure 4-.**
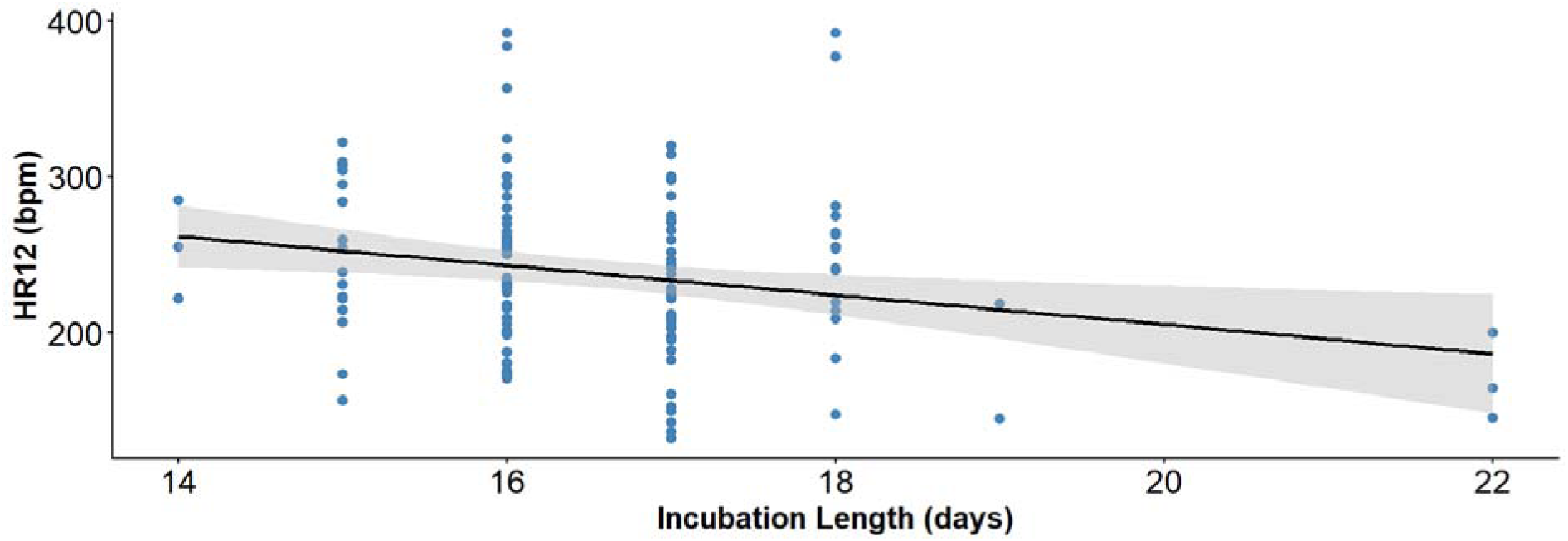
The correlation between incubation length (N = 134, 2018 = 27, 2019 = 18, 2023 = 89) on HR12 (heart rate at 12 of full-time incubation) in thorn tailed Rayadito in Navarino in 2018, 2019 & 2023. For statistical analyses, see table 2A.

## DISCUSSION

This study investigated whether climate predicts embryonic development in Thorn-tailed Rayadito embryos in Patagonia and if so, at what phase of development climate predict development. We did this at two study sites in Chile across different years which varies in climatic conditions. We measured embryonic development through measuring embryonic heart rate (HR12) just before hatching, which is agreed as a reliable and robust indicator for embryonic development (Sheldon et al., 2018 and Du et al., 2010).

This is the first study of its kind to the best of our knowledge to examine embryonic development in the field and to associate this to changing climatic conditions whilst highlighting specific days on which development is influenced. We found that minimum temperatures, wind speeds and year influenced embryonic development in Pucón. Lower minimum temperatures correlated with less embryonic development, extremes in wind speeds correlated with less embryonic growth and embryonic development on day 12 is getting lower each year. We find that rainfall, egg mass, clutch size, clutch attempt and incubation lengths have no impact on embryonic development in Pucón. Whilst in Navarino, only incubation lengths influence embryonic development with longer incubation lengths being associated with higher embryonic development. Climatic factors, year, clutch size and egg mass have no effect on embryonic development. This shows two contrasting reactions to the same selection pressures. Our main finding is that in Navarino, when embryonic development has been slow, incubation lengths are extended to mitigate any potential negative effects, whilst in Pucón, incubation lengths have not been lengthened enough to mitigate any negative effects.

### Location based effects

We attempted to combine data from both locations in our analysis but found conflicting results. For example, wind speeds have no effect on HR12 when data from both locations were combined. But when we analyse locations separately, wind speeds have a significant effect in Pucón but not in Navarino. Navarino had a dominant influence on the overall results when locations were combined. Analysing locations separately captures more detail on the specific effects of climate on different populations which are responding differently to the same selection pressures. This also highlights differences in the evolutionary trajectory of these separate populations as they are not moving in the same direction as they’re not being exposed to the same intensity of selection pressure. Therefore, analysing locations separately better answers our research questions. We have two separate but genetically similar Rayadito populations, one population (Pucón) has not shown adaptation to changing climatic conditions and the other population (Navarino) has shown adaptation to changing climatic conditions. There are several reasons why this difference could exist between the two populations. One reason is their relative isolation from each other. Evolving in separate locations means they respond differently to different selection pressures that have shaped their evolutionary trajectories. For example, average wind speeds are almost three times higher in Navarino (∼17 km/h) than in Pucón (∼6 km/h) because of the mountain ranges surrounding the site in Pucón (Table 1), so this population has not evolved to cope with high wind speeds, whilst the population in Navarino regularly encounters strong gusts and high average wind speeds. Additionally, HR12 is higher in the southern Navarino population compared to the northern Pucón. Given the consistent harsh winters in the south of Patagonia compared to the north this may be the reason why birds in this area are responding better to climate-based selection pressures, as they are better adapted to dealing with more variable climate.

### Drivers of embryonic development in Rayaditos

In the north of Patagonia (Pucón) varying climate generally has a negative effect on embryonic development. We found that low daily minimum temperatures and extremes in wind speeds correlate with low HR12 and indicate slow embryonic development. We found no evidence that the incubation period was prolonged to control for this, evidenced by the fact that incubation lengths did not predict HR12. However, it is possible that this will change over time. One reason why this has not occurred yet could be because climatic changes have occurred only very recently, and there may not be enough selection pressure for parents to respond to these changes. The alternative is that embryos and adults in Pucón are not responding to climate-based selection pressures and as a result, embryonic development will continue to decline.

Temperature is a well-documented predictor of embryonic development (Gilooly et al., 2002; Du et al., 2011; Sheldon et al., 2018). A specific study coherent with our findings on the relationship between climate and embryonic development is from Andrewartha, Tazawa and Burggren, (2011) who conducted lab experiments on chicken (*Gallus gallus domesticus)*, emu (*Dromaius novaehollandiae)* and duck (*Cairina moschata*) eggs where they exposed eggs to extreme low ambient temperatures and found that lower minimum temperatures (6-8 degrees) caused low embryonic heart rates near hatching. This aligns with our finding on minimum temperatures where lower daily minimum temperatures correlated with lower HR12, showing that lab-based studies can and do reflect field-based findings.

Wind speeds had a more complicated quadratic effect on HR12. During high wind speeds (>10km/h) around the time of clutch completion and the initiation of full-time incubation, HR12 was lower. Moderate wind speeds (∼4km/h) around the same time correlated with the highest HR12s and low wind speeds (<2km/h) correlated with low HR12. We are not sure why low and high wind speeds influenced embryonic development. In the case of high wind speeds, this is unlikely to be from a direct cooling effect as nest boxes generally protect from wind. This is more likely to be an indirect effect. One possibility is that high wind speeds may have a negative effect on the foraging ability of parents, resulting in more off bouts from the nest (Shoji et al., 2011) which can lead to lower incubation attendance and slower embryonic development (Ricklefs, Austin and Robinson, 2017). We equally do not understand why low wind speeds correlated with slower embryonic development. We can only speculate that low wind speeds had an interactive effect with other climatic variables that may have contributed to interfering with incubation patterns, similarly to the effect of high wind speeds. Current evidence suggests that wind speeds are due to increase significantly over the next century in northern Patagonia (Chemke, Ming and Yuval, 2022) and in the sub-Antarctic regions (Yu and Zhong, 2019) which are likely to have an impact on embryonic development as we have observed here. Incubation periods will need to be extended to produce high quality nestlings as seen in Navarino, or embryos will need to evolve to withstand more extreme climatic conditions (Du and Shine, 2022).

The year also showed a clear correlation on HR12 with embryonic HR12 being lower in 2018 compared to 2019, this is a decrease over time but having only two years of data makes it difficult to draw any strong conclusions. We would need more evidence to expand on the theory that embryonic development is becoming less over time, but we cannot ignore the trends we have found. This theory is supported by the strong evidence base that climate is becoming more varied across Patagonia (Pendlebury and Barnes-Keoghan, 2007; Yu and Zhong, 2019; Natalia et al., 2020; Chemke, Ming and Yuval, 2022) which as we have discovered here, can predict embryonic development. This is interesting because we find the population in southern Patagonia responds very differently to changing climatic conditions.

In Navarino, in the south of Patagonia, only incubation length predicted embryonic development. The reasons for this vary, but longer incubation periods give evidence to the theory that behaviourally, Rayaditos are adapting to changing climatic conditions in Navarino by prolonging incubation periods when conditions have been unfavourable. We cannot be sure if it is the adults or the embryos who are extending the incubation period. The theory that embryos dictate the incubation length was theorised by Du and Shine (2022), who propose that embryos can respond dynamically to their environment and adapt to changing conditions. Whereas previously, it was assumed that the parents are in control of when embryos hatch through controlling incubation patterns. In the case of the Rayadito, we cannot be sure if extending incubation patterns is a strategy by parents to ensure normal development or if it is a consequence of reduced embryonic development. As Du and Shine (2022) propose, it is more likely the embryos who decide when they hatch and parents just incubate until hatching. In any case, we have observed that when climate conditions are unfavourable in Navarino, the incubation period is prolonged. As incubation lengths were the only predictor of embryonic development, this clearly shows that Rayaditos avoid any negative climatic or other consequences by extending incubation periods. Although these trends are based on limited data (2-3 years) we cannot ignore the trend that incubation lengths are lengthening due to embryos taking longer to develop in Navarino. The short and long terms effects of this in Navarino are uncertain. However, in another study, we found that poorer climatic conditions during early incubation correlated with smaller hatchlings in Navarino (Badji-Churchill et al., 2026). Which overall, shows that longer incubation periods are leading to smaller hatchlings regardless of embryonic development, which is what other studies also find (Vedder et al., 2018; Stier, Metcalfe and Monaghan, 2020). However, it is also known that longer incubation periods produce smaller nestlings, which in Navarino, will make it more difficult for nestlings to survive their first winter, but smaller nestlings are also known to have longer telomere lengths which indicates longer life spans (Vedder et al., 2018; Stier, Metcalfe and Monaghan, 2020). These studies and our data combined reflect that although nestlings in Navarino may be at a short-term disadvantage by being born smaller (and therefore less likely to survive their first year), they may have long-term advantages through longer life spans.

### Fitness consequences of embryonic development

The consequences of lower HR12 will either result in underdeveloped nestlings that will struggle to thermoregulate and be of lower quality at hatching, or the incubation period will be prolonged, in which case nestlings may benefit from longer parental provisioning, as seen in other studies (Santos and Nakagawa., 2012; Klug and Bonsall, 2014; Vedder et al.,, 2018). We know that in Pucón, incubation lengths are not different between years, but HR12 is different between years (lower in 2019 than in 2018), which shows that parents are not incubating eggs for longer when embryos are underdeveloped. This is concerning as this is likely leading to poorer quality hatchlings. A future exploration of these findings should aim to assess what impact this has on nestlings during the rearing period and if this goes on to affect their survival rates or reproductive success, such a study could reveal if there is a trade-off during nestling development which corrects for this poor start in life. A meta-analysis of this by Krist (2011) revealed that poor egg quality overwhelmingly correlates with poorer quality nestlings and lower survival rates during the nestling phase. However, primary data is needed to establish a causal link between egg quality and nestling survival/reproduction in our study. We are lacking data of the full life trajectory of individuals; if lower quality eggs result in lower quality nestlings and if this in turn results in lower survival rates of nestlings, does this affect reproductive or survival rates in adults in the field? This would establish what factors are overall most important for a survival of a species and how selection pressures shape populations demographics under the destructive force of climate change.

In Navarino, we found that incubation lengths were shorter in 2019 compared to 2023, otherwise there was no difference between years, and there were no differences between years in HR12. Incubation lengths were longer when embryonic development was slower, which shows that when nestlings are underdeveloped, the incubation period was extended and this has a positive effect on embryonic development, as seen elsewhere (Santos & Nakagawa, 2012 & Vedder et al., 2018). Studies on wild wood ducks (*Aix sponsa*) found that colder incubation temperatures reduce, whereas slightly warmer incubation temperatures increase offspring cold tolerance (DuRant et al., 2012 and 2013). However, this temperature change in cold tolerance may not be that important as a study in common terns (*Sterna hirundo*) found that cooler incubation temperatures correlated with longer incubation periods and smaller nestlings, but this resulted in longer telomere lengths and in turn the potential for longer life spans (Vedder et al., 2018). This is important for our study as the consequences of slow embryonic development may not be only negative or neutral. In Navarino, where incubation lengths are longer, individuals may also develop longer telomere lengths and gain increased life spans because of extended incubation periods. So, although adverse climatic conditions during incubation may hinder embryonic development in the short-term, if incubation lengths are longer, it could provide long-term benefits to survival and reproduction.

### Conclusions and recommendations for future research

In conclusion, we found that the reaction of embryonic development to climatic conditions in the field is not homogenous. Embryos at high and low latitudes react differently to climatic conditions before and during embryonic development and the uniqueness of the climate in these two locations is driving these populations in different evolutionary directions. Climatic selection pressures are predicted to get drier, warmer, and windier with more extremes in daily minimum and maximum temperatures which will intensify climate as a selection pressure on embryonic development. Optimistically, embryos could react to these changes dynamically and adapt to their changed environments as discussed in (Du and Shine 2022 & Cones, Schneider & Westneat, 2024) or we may observe a continuing trade-off with longer incubation periods as observed in Navarino, or we may witness declining development as seen in Pucón. There is evidence to be optimistic that adults and/or embryos can react to unfavourable climatic conditions by extending incubation periods, which could lead to longer lived nestlings (Vedder et al, 2018; Stier, Metcalfe and Monaghan, 2020). Although, the impact of extended parental dependency would need to be closely studied. Another important investigation that should take place is egg swapping experiments between the two locations; this has been attempted in the past but the lack of available eggs and the timing of breeding between the two populations has not aligned. An experiment of this nature would reveal if embryonic adaptations were genetically or behaviourally driven, this would also help to answer if it is the embryos or the adults controlling the incubation length. These questions are essential to answer if we are to understand how these populations may change or adapt in the future to the irreversible and devastating climatic changes we are experiencing.

## Supporting information

Data and R scripts

