## Supplementary material for "Climatic conditions predict embryonic development in thorn tailed Rayadito (*Aphrastura spinicauda*)": Data and R scripts: Embryonic_Data_Modelling.R

```
library(climwin)
library(ggplot2) 
library(lme4)
library(DHARMa)
library(PerformanceAnalytics)
library(Hmisc)
library(missMDA)
library(FactoMineR)
library(factoextra)
library(mgcv)
library(dplyr)
library(emmeans)
library(lattice)
library(car)
library(lme4)

setwd("C:/Users/Owner/OneDrive/Documents/Rayadito PhD/Embryonic Development/CSV_Data")

ENV <- read.csv("ENV_DATA_ALL_1820.csv")
NVENV <- read.csv("ENV_DATA_NVRN_ALL.csv", sep=",")
PCENV <- read.csv("ENV_DATA_PCN_ALL.csv", sep=",")


EGG12 <- read.csv("EGG_DATA_CC12.csv", sep =",")
ALLDATA <- read.csv("ALL_EGG_DATA.csv", sep=",")

NVEGG12 <- read.csv("EGG_DATA_CC12_NVRN.csv", sep=",")
PCEGG12 <- read.csv("EGG_DATA_CC12_PCN.csv", sep=",")


NVENV$LOCATION <- as.factor(NVENV$LOCATION)
NVENV$YEAR <- as.factor(NVENV$YEAR)
NVENV$MONTH <- as.factor(NVENV$MONTH)
NVENV$NEWDATE <- as.Date(NVENV$DATE,
                         format = "%d/%m/%Y")
NVENV$MONTHYEAR <- interaction(NVENV$MONTH, NVENV$YEAR)

PCENV$LOCATION <- as.factor(PCENV$LOCATION)
PCENV$YEAR <- as.factor(PCENV$YEAR)
PCENV$MONTH <- as.factor(PCENV$MONTH)
PCENV$NEWDATE <- as.Date(PCENV$DATE,
                         format = "%d/%m/%Y")
PCENV$MONTHYEAR <- interaction(PCENV$MONTH, PCENV$YEAR)

EGG12$LOCATION <- as.factor(EGG12$LOCATION)
EGG12$YEAR <- as.factor(EGG12$YEAR)
EGG12$NB <- as.numeric(EGG12$NB)
EGG12$Location_code <- as.factor(EGG12$Location_code)
EGG12$NEST_ID_LOCATION <- as.factor(EGG12$NEST_ID_LOCATION)
EGG12$EggID <- as.factor(EGG12$EggID)

EGG12$NEWDATE <- as.Date(EGG12$Measure_date,
                         format = "%d/%m/%Y")

EGG12$NEST_ID_YEAR <- interaction(EGG12$NB, EGG12$YEAR)
EGG12$GROUP <- interaction(EGG12$LOCATION, EGG12$YEAR)

EGG12$Egg_no <- as.factor(EGG12$Egg_no)
EGG12$CC_age <-as.factor(EGG12$CC_age)
EGG12$Nest_temp <- as.numeric(EGG12$Nest_temp)
EGG12$Egg_temp <- as.numeric(EGG12$Egg_temp)
EGG12$HR <- as.numeric(EGG12$HR)
EGG12$Mass <- as.numeric(EGG12$Mass)
EGG12$Time <- as.POSIXlt(as.character(EGG12$Time), format="%H:%M:%S")


ENV$LOCATION <- as.factor(ENV$LOCATION)
ENV$Av_temp <- as.numeric(ENV$Av_temp)
ENV$Av_precip <- as.numeric(ENV$Av_precip)
ENV$Av_rh <- as.numeric(ENV$Av_rh)
ENV$Av_ws_12 <- as.numeric(ENV$Av_ws_12)
ENV$Acc_mill <- as.numeric(ENV$Acc_mill)
ENV$NEWDATE <- as.Date(ENV$DATE,
                       format = "%d/%m/%Y")

str(ALLDATA)

ALLDATA$LOCATION <- as.factor(ALLDATA$LOCATION)
ALLDATA$YEAR <- as.factor(ALLDATA$YEAR)
ALLDATA$NEST_BOX <- as.numeric(ALLDATA$NEST_BOX)
ALLDATA$EGG_ID <- as.factor(ALLDATA$EGG_ID)
ALLDATA$NEW_CC_DATE <- as.Date(ALLDATA$CC_DATE,
                               format = "%d/%m/%Y")
ALLDATA$GROUP <- interaction(ALLDATA$LOCATION, ALLDATA$YEAR)

str(NVEGG12)

NVEGG12$LOCATION <- as.factor(NVEGG12$LOCATION)
NVEGG12$YEAR <- as.factor(NVEGG12$YEAR)
NVEGG12$NEST_BOX <- as.factor(NVEGG12$NEST_BOX)
NVEGG12$EGG_ID <- as.factor(NVEGG12$EGG_ID)
NVEGG12$INCUBATION_LENGTH <- as.integer(NVEGG12$INCUBATION_LENGTH)
NVEGG12$EGG_NO <- as.factor(NVEGG12$EGG_NO)
NVEGG12$ORDER <- as.factor(NVEGG12$ORDER)
NVEGG12$CLUTCH <- as.numeric(NVEGG12$CLUTCH)
NVEGG12$CLUTCH_ATTEMPT <- as.factor(NVEGG12$CLUTCH_ATTEMPT)
NVEGG12$NEWDATE <- as.Date(NVEGG12$MEASURE_DATE,
                           format = "%d/%m/%Y")
NVEGG12$CLUTCHf <- as.factor(NVEGG12$CLUTCH)
NVEGG12$NB_YEAR <- paste(NVEGG12$NEST_BOX, NVEGG12$YEAR, sep = "_")
NVEGG12$NB_YEAR <- as.factor(NVEGG12$NB_YEAR)

PCEGG12$LOCATION <- as.factor(PCEGG12$LOCATION)
PCEGG12$YEAR <- as.factor(PCEGG12$YEAR)
PCEGG12$NEST_BOX <- as.factor(PCEGG12$NEST_BOX)
PCEGG12$EGG_ID <- as.factor(PCEGG12$EGG_ID)
PCEGG12$EGG_NO <- as.factor(PCEGG12$EGG_NO)
PCEGG12$ORDER <- as.factor(PCEGG12$ORDER)
PCEGG12$CLUTCH <- as.numeric(PCEGG12$CLUTCH)
PCEGG12$CLUTCH_ATTEMPT <- as.factor(PCEGG12$CLUTCH_ATTEMPT)
PCEGG12$NEWDATE <- as.Date(PCEGG12$MEASURE_DATE,
                           format = "%d/%m/%Y")
PCEGG12$CLUTCHf <- as.factor(PCEGG12$CLUTCH)


PCEGG12$NB_YEAR <- paste(PCEGG12$NEST_BOX, PCEGG12$YEAR, sep = "_")
PCEGG12$NB_YEAR <- as.factor(PCEGG12$NB_YEAR)


#
##           FULL ANALYSIS OF EMBRYONIC DEVELOPMENT

# Locations analysed together

HRD <- lm(HR ~ LOCATION, data=EGG12) # Pucon has lower HRs at CC+12 - another reason to analyse separately
summary(HRD)
boxplot(HR ~ LOCATION,data=EGG12)

tapply(PCEGG12$NEST_BOX[PCEGG12$YEAR %in% c(2018, 2019)],
       PCEGG12$YEAR[PCEGG12$YEAR %in% c(2018, 2019)],
       function(x) length(unique(x)))

with(PCENV[PCENV$YEAR %in% c(1819, 1920), ],
     tapply(Acc_mill, YEAR, mean, na.rm = TRUE))


with(NVENV[NVENV$YEAR %in% c(1819, 1920, 2023), ],
     tapply(Acc_mill, YEAR, mean, na.rm = TRUE))

plot(NVEGG12$INCUBATION_LENGTH)
plot(PCEGG12$INCUBATION_LENGTH)

mean(NVEGG12$INCUBATION_LENGTH, na.rm = TRUE)
mean(PCEGG12$INCUBATION_LENGTH, na.rm = TRUE)


t.test(PCEGG12$INCUBATION_LENGTH,NVEGG12$INCUBATION_LENGTH) # Not different
t.test(MASS ~ LOCATION, data = ALLDATA) # Bigger in Navarino
boxplot(MASS ~ LOCATION, 
        data = ALLDATA,
        xlab = "Location",
        ylab = "Mass",
        main = "Boxplot of Mass by Location",
        col = "lightgray")

MAD <- lm(MASS ~ CC_AGE, data=ALLDATA)
summary(MAD)
plot(MASS ~ CC_AGE, data=ALLDATA)

NVEGG12 %>%
  filter(YEAR %in% c(2018, 2019, 2023)) %>%
  group_by(YEAR) %>%
  summarise(
    n = n(),
    
    mean_INCUBATION_LENGTH = mean(INCUBATION_LENGTH, na.rm = TRUE),
    min_INCUBATION_LENGTH  = min(INCUBATION_LENGTH, na.rm = TRUE),
    max_INCUBATION_LENGTH  = max(INCUBATION_LENGTH, na.rm = TRUE),
    
    mean_HR = mean(HR, na.rm = TRUE),
    min_HR  = min(HR, na.rm = TRUE),
    max_HR  = max(HR, na.rm = TRUE)
  )

PCEGG12 %>%
  filter(YEAR %in% c(2018, 2019, 2023)) %>%
  group_by(YEAR) %>%
  summarise(
    n = n(),
    
    mean_INCUBATION_LENGTH = mean(INCUBATION_LENGTH, na.rm = TRUE),
    min_INCUBATION_LENGTH  = min(INCUBATION_LENGTH, na.rm = TRUE),
    max_INCUBATION_LENGTH  = max(INCUBATION_LENGTH, na.rm = TRUE),
    
    mean_HR = mean(HR, na.rm = TRUE),
    min_HR  = min(HR, na.rm = TRUE),
    max_HR  = max(HR, na.rm = TRUE)
  )

NVEGG12 %>%
  filter(YEAR %in% c(2018, 2019, 2023)) %>%
  group_by(YEAR) %>%
  summarise(
    n = n(),
    
    mean_HR = mean(HR, na.rm = TRUE),
    min_HR  = min(HR, na.rm = TRUE),
    max_HR  = max(HR, na.rm = TRUE),
    
    mean_HR = mean(HR, na.rm = TRUE),
    min_HR  = min(HR, na.rm = TRUE),
    max_HR  = max(HR, na.rm = TRUE)
  )

PCEGG12 %>%
  filter(YEAR %in% c(2018, 2019, 2023)) %>%
  group_by(YEAR) %>%
  summarise(
    n = n(),
    
    mean_HR = mean(HR, na.rm = TRUE),
    min_HR  = min(HR, na.rm = TRUE),
    max_HR  = max(HR, na.rm = TRUE),
    
    mean_HR = mean(HR, na.rm = TRUE),
    min_HR  = min(HR, na.rm = TRUE),
    max_HR  = max(HR, na.rm = TRUE)
  )


#######
#               Pucon
######

MD2 <- lm(HR ~ EGG_NO, data=PCEGG12)  # No differences
summary(MD2)
lsmeans(MD2, pairwise~EGG_NO)

ORDPC <- lm(HR ~ ORDER, data=PCEGG12) # No difference in HR egg order - eggs measured last did not have lower HRs
summary(ORDPC)

CAM <- lm(HR ~ CLUTCH_ATTEMPT, data = PCEGG12) # No differences between 1st or 2nd clutch attempt
summary(CAM)
 
CAM1 <- lm(HR ~ CLUTCHf, data = PCEGG12) # No difference in HR between clutch sizes
summary(CAM1)
lsmeans(CAM1, pairwise ~ CLUTCHf, adjust = "tukey")
table(PCEGG12$CLUTCHf)
table(NVEGG12$CLUTCHf)

CAM2 <-lm(HR~CLUTCH, data=PCEGG12)
summary(CAM2)
plot(HR~CLUTCH, data=PCEGG12) # looks linear with larger clutches = Higher HRs

HRINCP <- lm(HR ~ INCUBATION_LENGTH, data=PCEGG12) # Longer incubation = lower HRs on CC+12
summary(HRINCP)

PCEY <- lm(HR ~ YEAR, data=PCEGG12) # HR is lower in 2019 than in 2018
summary(PCEY)
plot(HR~YEAR, data=PCEGG12)

ILY <- lm(INCUBATION_LENGTH ~ YEAR, data=PCEGG12) # No difference
summary(ILY)
plot(INCUBATION_LENGTH~YEAR, data=PCEGG12)


# Full models

str(PCEGG12)
colSums(is.na(PCEGG12))
PCEGG12 <- PCEGG12[!is.na(PCEGG12$INCUBATION_LENGTH), ] # two NAs in Incubation length of two eggs that were eaten between CC12-Hatching

FULLMODEL2 <- lmer(log10(HR) ~ MIN_TEMP_LIN + RAIN_LIN + WIND_LIN + WIND_QUAD + YEAR + MASS + CLUTCH + INCUBATION_LENGTH + CLUTCH_ATTEMPT + (1|NB_YEAR), data=PCEGG12)
summary(FULLMODEL2)
vif(FULLMODEL2) 
drop1(FULLMODEL2, test="Chisq");
Sim=simulateResiduals(FULLMODEL2, re.form = ~0, plot = T)  # Residuals fine
testDispersion(Sim)


# Model run with "outliers" removed

PCEGG12_RMO <- PCEGG12[-c(), ]

FULLMODEL3 <- lmer(log10(HR) ~ MIN_TEMP_LIN + RAIN_LIN + WIND_LIN + WIND_QUAD + YEAR + MASS + CLUTCH + INCUBATION_LENGTH + CLUTCH_ATTEMPT + (1|NEST_BOX), data=PCEGG12_RMO)
summary(FULLMODEL3)
vif(FULLMODEL3) 
drop1(FULLMODEL3, test="Chisq");
Sim=simulateResiduals(FULLMODEL3, re.form = ~0, plot = T)  # Residuals fine
testDispersion(Sim)

Anova(FULLMODEL3, type = "III")


#######
########          OUTLIER TEST

z_MIN_TEMP <- scale(PCEGG12$MIN_TEMP_LIN)
z_WIND_LIN <- scale(PCEGG12$WIND_LIN)
z_WIND_QUAD <- scale(PCEGG12$WIND_QUAD)

which(abs(z_MIN_TEMP) > 3)
which(abs(z_WIND_LIN) > 3)
which(abs(z_WIND_QUAD) > 3)

outlierTest <- which(abs(Sim$scaledResiduals) > 3)
outlierTest

# PCN YEAR AND HR BOXPLOT

ggplot(PCEGG12, aes(x = YEAR, y = HR, fill = YEAR)) +
  geom_boxplot(color = "black", width = 0.6, outlier.color = "red", outlier.shape = 16) +
  scale_fill_manual(values = c("2018" = "#9CAF88", "2019" = "#FFF9C4")) +  # Sage green and light yellow
  theme_minimal(base_size = 14) +
  labs(
    title = "Heart Rates by Year",
    x = "Year",
    y = "Heart Rates (bpm)"
  ) +
  theme(
    plot.title = element_text(hjust = 0.5, face = "bold"),
    axis.title = element_text(face = "bold"),
    axis.text = element_text(color = "black", size = 16),
    axis.line = element_line(color = "black"),
    axis.ticks = element_line(color = "black"),
    panel.grid.major = element_blank(),
    panel.grid.minor = element_blank(),
    panel.border = element_blank(),
    legend.position = "none"
  )


##      Wind speeds Pucon grouped by year

ggplot(PCEGG12, aes(x = WIND_LIN, y = HR, color = as.factor(YEAR))) +
  geom_point(alpha = 0.6, size = 3) +
  geom_smooth(data = subset(PCEGG12, YEAR %in% c(2018, 2019)),
              aes(group = YEAR),
              method = "lm",
              formula = y ~ x + I(x^2),
              se = FALSE,
              color = "black",
              linewidth = 2.2) +
  geom_smooth(data = subset(PCEGG12, YEAR %in% c(2018, 2019)),
              aes(group = YEAR),
              method = "lm",
              formula = y ~ x + I(x^2),
              se = TRUE,
              linewidth = 1.5) +
  scale_color_manual(values = c("2018" = "#1f78b4",  
                                "2019" = "#e31a1c"), 
                     name = "Year") +
  labs(
    x = expression("Wind speed (km/h) 12-13 days before HR12 measured"),
    y = "HR12 (bpm)",
    color = "Year"
  ) +
  theme_minimal(base_size = 18) +
  theme(
    plot.title = element_text(hjust = 0.5, face = "bold", size = 20),
    axis.title = element_text(face = "bold", size = 18),
    axis.text = element_text(color = "black", size = 16),
    axis.line = element_line(color = "black"),
    axis.ticks = element_line(color = "black"),
    panel.grid.major = element_blank(),
    panel.grid.minor = element_blank(),
    panel.border = element_blank(),
    legend.position = "right"
  )


# Plot of Min_temp and HR in Pucon grouped by year

ggplot(PCEGG12, aes(x = MIN_TEMP_LIN, y = HR, color = as.factor(YEAR))) +
  geom_point(size = 3, alpha = 0.7) +
  geom_smooth(data = subset(PCEGG12, YEAR %in% c(2018, 2019)),
              aes(group = YEAR),
              method = "lm",
              se = FALSE,
              color = "black",
              linewidth = 2.2) +
  geom_smooth(data = subset(PCEGG12, YEAR %in% c(2018, 2019)),
              aes(group = YEAR),
              method = "lm",
              se = TRUE,
              linewidth = 1.5) +
  scale_color_manual(values = c("2018" = "#1f78b4",  
                                "2019" = "#e31a1c"),
                     name = "Year") +
  labs(
    x = "Minimum temperature (°C) 6-13 days before HR12 measured",
    y = "HR12 (bpm)",
    color = "Year"
  ) +
  theme_minimal(base_size = 18) +
  theme(
    plot.title = element_text(hjust = 0.5, face = "bold", size = 20),
    axis.title = element_text(face = "bold", size = 18),
    axis.text = element_text(color = "black", size = 16),
    axis.line = element_line(color = "black"),
    axis.ticks = element_line(color = "black"),
    panel.grid.major = element_blank(),
    panel.grid.minor = element_blank(),
    panel.border = element_blank(),
    legend.position = "right"
  )

#######
#               Navarino
######
 
# No weather variables were found to effect embryonic HRs in Navarino

str(NVEGG12)

MD1 <- lm(HR ~ EGG_NO, data=NVEGG12)  # No differences
summary(MD1)
lsmeans(MD1, pairwise~EGG_NO)

MD2 <- lm(HR ~ ORDER, data=NVEGG12)  # Order did matter and cooling did have an effect here so first measured egg had highest HR.
summary(MD2)
lsmeans(MD2, pairwise~ORDER)
plot(HR~ORDER, data=NVEGG12)

HRINC <- lm(HR ~ INCUBATION_LENGTH, data=NVEGG12) # Longer incubation lengths have lower HRs at CC+12
summary(HRINC)
plot(NVEGG12$HR,NVEGG12$INCUBATION_LENGTH) # Longer incubation lengths have lower HRs at CC+12

NVY <- lm(HR ~ YEAR, data=NVEGG12) # 2023 had lower HR than 2018 but no different to 2019
summary(NVY)
plot(HR~YEAR, data=NVEGG12) # 2018 has highest HR overall adn 2023 = lowest 
lsmeans(NVY, pairwise~YEAR) 

ILNVY <- lm(INCUBATION_LENGTH~YEAR, data=NVEGG12)
summary(ILNVY) # No difference in incubation length between years OVERALL
lsmeans(ILNVY, pairwise~YEAR) # However, 2019 has significantly lower incubation lengths than 2023


ggplot(NVEGG12, aes(x = factor(YEAR), y = INCUBATION_LENGTH)) +
  geom_boxplot(fill = "#A8D5BA", color = "black", width = 0.6, outlier.color = "red", outlier.shape = 16) +
  theme_minimal(base_size = 14) +
  labs(
    x = "Year",
    y = "Incubation Length (days)"
  ) +
  theme(
    plot.title = element_text(hjust = 0.5, face = "bold", size = 16),
    axis.title = element_text(face = "bold"),
    axis.text = element_text(color = "black", size = 16),
    axis.line = element_line(color = "black"),
    axis.ticks = element_line(color = "black"),
    panel.grid.major = element_blank(),
    panel.grid.minor = element_blank(),
    panel.border = element_blank()
  )


# Full model

colSums(is.na(NVEGG12))
NVEGG12 <- NVEGG12[!is.na(NVEGG12$INCUBATION_LENGTH), ] # 7 Incubation length NAs so need to remove to include
table(NVEGG12$YEAR)

FULLNVM2 <- lmer(log10(HR) ~ YEAR + CLUTCH + MASS + INCUBATION_LENGTH + (1|NB_YEAR), data=NVEGG12)
summary(FULLNVM2)
vif(FULLNVM2) 
drop1(FULLNVM2, test="Chisq");
Sim=simulateResiduals(FULLNVM2, re.form = ~0, plot = T)  # Residuals fine
testDispersion(Sim)

# Plot of incubation length and HRs

ggplot(NVEGG12, aes(x = INCUBATION_LENGTH, y = HR)) +
  geom_point(color = "steelblue", size = 2) +
  geom_smooth(method = "lm", color = "black", fill = "grey70", se = TRUE) +
  labs(
    x = "Incubation Length (days)",
    y = "HR12 (bpm)"
  ) +
  theme_classic() +
  theme(
    plot.title = element_text(size = 16, face = "bold", hjust = 0.5),
    axis.text = element_text(size = 16, color = "black"),
    axis.title = element_text(size = 15, face = "bold", color = "black"),
    axis.line = element_line(color = "black"),
    panel.grid.major = element_blank(),
    panel.grid.minor = element_blank()
  )


# Plot of egg order and HR


NVEGG12_clean <- subset(NVEGG12, !is.na(ORDER))


ggplot(NVEGG12_clean, aes(x = ORDER, y = HR, fill = as.factor(ORDER))) +
  geom_boxplot(color = "black", width = 0.6, outlier.color = "red", outlier.shape = 16) +
  scale_fill_manual(values = c(
    "#A8DADC",
    "#457B9D",  
    "#F4A261",  
    "#E76F51",  
    "#2A9D8F",  
    "#B5838D"   
  )) +
  theme_minimal(base_size = 14) +
  labs(
    title = "Heart Rate by Measuring Order",
    x = "Order Measured",
    y = "Heart Rates (bpm)"
  ) +
  theme(
    plot.title = element_text(hjust = 0.5, face = "bold"),
    axis.title = element_text(face = "bold"),
    axis.text = element_text(color = "black", size = 16),
    axis.line = element_line(color = "black"),
    axis.ticks = element_line(color = "black"),
    panel.grid.major = element_blank(),
    panel.grid.minor = element_blank(),
    panel.border = element_blank(),
    legend.position = "none"
  )
```
