## Supplementary material for "Climatic conditions predict embryonic development in thorn tailed Rayadito (*Aphrastura spinicauda*)": Data and R scripts: Embryonic_Dev_D12Clim.R

```
############################################################
############################################################
############################################################

############      EGG HEART RATES AND CLIMATE NVRN    ###########

############################################################
############################################################


hist(NVEGG12$HR)
hist(log10(NVEGG12$HR))
NVEGG12=subset(NVEGG12, !is.na(HR) & HR!=0)

#     AVERAGE TEMPS AND HR NVRN


HRAVTEMPEGGNVRN <- slidingwin(xvar = list(Temp = NVENV$Av_temp),
                     exclude = NA,
                     cdate = NVENV$NEWDATE,
                     bdate = NVEGG12$NEWDATE,
                     baseline = lmer(log10(HR) ~ YEAR + (1|NEST_BOX), data = NVEGG12),
                     cinterval = "day",
                     range = c(20, 0),
                     type = "relative",
                     stat = c("mean"),
                     func = c("lin","quad"))

HRAVTEMPEGGNVRN$combos
summary(HRAVTEMPEGGNVRN[[1]]$BestModel)


## test if nestbox is an important random effect
lm1=lm(yvar~factor(YEAR)+climate, data=HRAVTEMPEGGNVRN[[1]]$BestModelData)
lmer1=lmer(yvar~factor(YEAR)+climate+(1|NEST_BOX), data=HRAVTEMPEGGNVRN[[1]]$BestModelData)
anova(lmer1,lm1)

lm1=lm(yvar~factor(YEAR)+climate, data=HRAVTEMPEGGNVRN[[2]]$BestModelData)
lmer1=lmer(yvar~factor(YEAR)+climate+(1|NEST_BOX), data=HRAVTEMPEGGNVRN[[2]]$BestModelData)
anova(lmer1,lm1)
# Nest box is important and model is better with it

plotwin(dataset=HRAVTEMPEGGNVRN[[1]]$Dataset)
plot(HRAVTEMPEGGNVRN[[1]]$BestModelData)

Data=HRAVTEMPEGGNVRN[[1]]$BestModelData

plot(Data$yvar~Data$climate)

Egg2 <- HRAVTEMPEGGNVRN[[1]]$Dataset
summary(Egg2)


RandEgg <- randwin(repeats = 1000, 
                   window = "sliding", 
                   xvar=list(Temp = NVENV$Av_temp), 
                   cdate=NVENV$NEWDATE,
                   bdate = NVEGG12$NEWDATE,
                   baseline= lmer(log10(HR) ~ YEAR + (1|NEST_BOX), data = NVEGG12), 
                   stat= "mean", 
                   range=c(20,0),
                   func="lin",
                   type ="relative",
                   cinterval = "day")


plotall(dataset = HRAVTEMPEGGNVRN[[1]]$Dataset, datasetrand = RandEgg[[1]],
        cw1 = 0.95, cw2 = 0.5, cw3 = 0.25,
        title = NULL, arrow = FALSE)

res_simlog <- DHARMa::simulateResiduals(HRAVTEMPEGGNVRN[[1]]$BestModel, re.form = ~ (1|NEST_BOX))
plot(res_simlog) 

####
#             MAXIMUM TEMPERATURE AND HR NVRN
####

NVEGG12$HR
hist(NVEGG12$HR)

HRMAXTEMPEGGNVRN <- slidingwin(xvar = list(Temp = NVENV$Max_Temp),
                              exclude = NA,
                              cdate = NVENV$NEWDATE,
                              bdate = NVEGG12$NEWDATE,
                              baseline = lmer(log10(HR) ~ YEAR + (1|NEST_BOX), data = NVEGG12),
                              cinterval = "day",
                              range = c(20, 0),
                              type = "relative",
                              stat = c("mean"),
                              func = c("lin","quad"))

HRMAXTEMPEGGNVRN$combos
summary(HRMAXTEMPEGGNVRN[[1]]$BestModel)

lm1=lm(yvar~factor(YEAR)+climate+I(climate^2) , data=HRMAXTEMPEGGNVRN[[2]]$BestModelData)
lmer1=lmer(yvar~factor(YEAR)+climate+I(climate^2)+(1|NEST_BOX), data=HRMAXTEMPEGGNVRN[[2]]$BestModelData)
anova(lmer1,lm1)

plotwin(dataset=HRMAXTEMPEGGNVRN[[1]]$Dataset)
plot(HRMAXTEMPEGGNVRN[[1]]$BestModelData)

Egg2 <- HRMAXTEMPEGGNVRN[[1]]$Dataset
summary(Egg2)


RandEgg <- randwin(repeats = 1000, 
                   window = "sliding", 
                   xvar=list(Temp = NVENV$Max_Temp), 
                   cdate=NVENV$NEWDATE,
                   bdate = NVEGG12$NEWDATE,
                   baseline= lmer(log10(HR) ~ YEAR + (1|NEST_BOX), data = NVEGG12), 
                   stat= "mean", 
                   range=c(20,0), 
                   func="lin",
                   type ="relative",
                   cinterval = "day")

plotall(dataset = HRMAXTEMPEGGNVRN[[1]]$Dataset, datasetrand = RandEgg[[1]],
        cw1 = 0.95, cw2 = 0.5, cw3 = 0.25,
        title = NULL, arrow = FALSE)

res_sim <- DHARMa::simulateResiduals(HRMAXTEMPEGGNVRN[[1]]$BestModel, re.form = ~ (1|NEST_BOX))
plot(res_sim)


####
#             MINIMUM TEMPERATURE AND HR NVRN
####

HRMINTEMPEGGNVRN <- slidingwin(xvar = list(Temp = NVENV$Min_Temp),
                               exclude = NA,
                               cdate = NVENV$NEWDATE,
                               bdate = NVEGG12$NEWDATE,
                               baseline = lmer(log10(HR) ~ YEAR + (1|NEST_BOX), data = NVEGG12),
                               cinterval = "day",
                               range = c(20, 0),
                               type = "relative",
                               stat = c("mean"),
                               func = c("lin","quad"))

HRMINTEMPEGGNVRN$combos
summary(HRMINTEMPEGGNVRN[[1]]$BestModel)

lm1=lm(yvar~factor(YEAR)+climate, data=HRMINTEMPEGGNVRN[[1]]$BestModelData)
lmer1=lmer(yvar~factor(YEAR)+climate+(1|NEST_BOX), data=HRMINTEMPEGGNVRN[[1]]$BestModelData)
anova(lmer1,lm1)


plotwin(dataset=HRMINTEMPEGGNVRN[[1]]$Dataset)
plot(HRMINTEMPEGGNVRN[[1]]$BestModelData)

Egg2 <- HRMINTEMPEGGNVRN[[1]]$Dataset
summary(Egg2)

RandEgg <- randwin(repeats = 1000, 
                   window = "sliding", 
                   xvar=list(Temp = NVENV$Min_Temp), 
                   cdate=NVENV$NEWDATE,
                   bdate = NVEGG12$NEWDATE,
                   baseline= lmer(log(HR) ~ YEAR + (1|NEST_BOX), data = NVEGG12), 
                   stat= "mean", 
                   range=c(20,0), 
                   func="lin",
                   type ="relative", 
                   cinterval = "day")

plotall(dataset = HRMINTEMPEGGNVRN[[1]]$Dataset, datasetrand = RandEgg[[1]],
        cw1 = 0.95, cw2 = 0.5, cw3 = 0.25,
        title = NULL, arrow = FALSE)

res_sim <- DHARMa::simulateResiduals(HRMINTEMPEGGNVRN[[1]]$BestModel, re.form = ~ (1|NEST_BOX))
plot(res_sim)

##############################################################

##            HR AND RAINFALL IN NVRN

##############################################################


HRRAINNVRN <- slidingwin(xvar = list(Rain = NVENV$Acc_mill),
                       exclude = NA,
                       cdate = NVENV$NEWDATE,
                       bdate = NVEGG12$NEWDATE,
                       baseline = lmer(log10(HR) ~ YEAR + (1|NEST_BOX), data = NVEGG12),
                       cinterval = "day",
                       range = c(20, 0),
                       type = "relative",
                       stat = c("mean"),
                       func = c("lin","quad"))

HRRAINNVRN$combos
summary(HRRAINNVRN[[1]]$BestModel)

lm1=lm(yvar~factor(YEAR)+climate+I(climate^2), data=HRRAINNVRN[[1]]$BestModelData)
lmer1=lmer(yvar~factor(YEAR)+climate+I(climate^2)+(1|NEST_BOX), data=HRRAINNVRN[[1]]$BestModelData)
anova(lmer1,lm1)


plotwin(dataset=HRRAINNVRN[[1]]$Dataset)
plot(HRRAINNVRN[[1]]$BestModelData)

Egg3 <- HRRAINNVRN[[1]]$Dataset
summary(Egg3)

RandEgg1 <- randwin(repeats = 1000, 
                   window = "sliding", 
                   xvar=list(Rain = NVENV$Acc_mill), 
                   cdate=NVENV$NEWDATE,
                   bdate = NVEGG12$NEWDATE,
                   baseline= lmer(log10(HR) ~ YEAR + (1|NEST_BOX), data = NVEGG12), 
                   stat= "mean", 
                   range=c(20,0), 
                   func="lin",
                   type ="relative", 
                   cinterval = "day")

plotall(dataset = HRRAINNVRN[[1]]$Dataset, datasetrand = RandEgg1[[1]],
        cw1 = 0.95, cw2 = 0.5, cw3 = 0.25,
        title = NULL, arrow = FALSE)

res_sim <- DHARMa::simulateResiduals(HRRAINNVRN[[1]]$BestModel, re.form = ~ (1|NEST_BOX))
plot(res_sim)


################################################################

########            HR AND WIND SPEED NVRN

################################################################

HRWINDNVRN <- slidingwin(xvar = list(Wind = NVENV$Av_ws_12),
                       exclude = NA,
                       cdate = NVENV$NEWDATE,
                       bdate = NVEGG12$NEWDATE,
                       baseline = lmer(log10(HR) ~ YEAR + (1|NEST_BOX), data = NVEGG12),
                       cinterval = "day",
                       range = c(20, 0),
                       type = "relative",
                       stat = c("mean"),
                       func = c("lin","quad"))

HRWINDNVRN$combos
summary(HRWINDNVRN[[1]]$BestModel)

lm1=lm(yvar~factor(YEAR)+climate, data=HRWINDNVRN[[1]]$BestModelData)
lmer1=lmer(yvar~factor(YEAR)+climate+(1|NEST_BOX), data=HRWINDNVRN[[1]]$BestModelData)
anova(lmer1,lm1)

plotwin(dataset=HRWINDNVRN[[1]]$Dataset)
plot(HRWINDNVRN[[1]]$BestModelData)

Egg4 <- HRWINDNVRN[[1]]$Dataset
summary(Egg4)

RandEgg2 <- randwin(repeats = 1000, 
                    window = "sliding", 
                    xvar=list(Wind = NVENV$Av_ws_12), 
                    cdate=NVENV$NEWDATE,
                    bdate = NVEGG12$NEWDATE,
                    baseline= lmer(log10(HR) ~ YEAR + (1|NEST_BOX), data = NVEGG12), 
                    stat= "mean", 
                    range=c(20,0),
                    func="lin",
                    type ="relative", 
                    cinterval = "day")

plotall(dataset = HRWINDNVRN[[1]]$Dataset, datasetrand = RandEgg2[[1]],
        cw1 = 0.95, cw2 = 0.5, cw3 = 0.25,
        title = NULL, arrow = FALSE)

res_sim <- DHARMa::simulateResiduals(HRWINDNVRN[[1]]$BestModel, re.form = ~ (1|NEST_BOX))
plot(res_sim)

############################################################
############################################################
############################################################

############      EGG HEART RATES AND CLIMATE PUCON    ###########

############################################################
############################################################

#     AVERAGE TEMP AND HR PCN

hist(log10(PCEGG12$HR))

HRAVTEMPEGGPC <- slidingwin(xvar = list(Temp = PCENV$Av_temp),
                              exclude = NA,
                              cdate = PCENV$NEWDATE,
                              bdate = PCEGG12$NEWDATE,
                              baseline = lmer(log10(HR) ~ YEAR + (1|NEST_BOX), data = PCEGG12),
                              cinterval = "day",
                              range = c(20, 0),
                              type = "relative",
                              stat = c("mean"),
                              func = c("lin","quad"))

HRAVTEMPEGGPC$combos

PCAVTEMPDATA=HRAVTEMPEGGPC[[1]]$BestModelData

PCEGG12$AvTemp=PCENV$Av_temp[match(PCEGG12$NEWDATE,PCENV$NEWDATE)]

plot(PCAVTEMPDATA$yvar~PCAVTEMPDATA$climate)
abline(lm(PCAVTEMPDATA$yvar~PCAVTEMPDATA$climate))
summary(HRAVTEMPEGGPC[[1]]$BestModel)

lm1=lm(yvar~factor(YEAR)+climate, data=HRAVTEMPEGGPC[[1]]$BestModelData)
lmer1=lmer(yvar~factor(YEAR)+climate+(1|NEST_BOX), data=HRAVTEMPEGGPC[[1]]$BestModelData)
anova(lmer1,lm1)

plotwin(dataset=HRAVTEMPEGGPC[[1]]$Dataset)
plot(HRAVTEMPEGGPC[[1]]$BestModelData)

Egg2 <- HRAVTEMPEGGPC[[1]]$Dataset
summary(Egg2)

RandEgg <- randwin(repeats = 1000, 
                   window = "sliding", 
                   xvar=list(Temp = PCENV$Av_temp), 
                   cdate=PCENV$NEWDATE,
                   bdate = PCEGG12$NEWDATE,
                   baseline= lmer(log10(HR) ~ YEAR + (1|NEST_BOX), data = PCEGG12), 
                   stat= "mean", 
                   range=c(20,0), 
                   func="lin",
                   type ="relative", 
                   cinterval = "day")

plotall(dataset = HRAVTEMPEGGPC[[1]]$Dataset, datasetrand = RandEgg[[1]],
        cw1 = 0.95, cw2 = 0.5, cw3 = 0.25,
        title = NULL, arrow = FALSE)

res_sim <- DHARMa::simulateResiduals(HRAVTEMPEGGPC[[1]]$BestModel, re.form = ~ (1|NEST_BOX))
plot(res_sim)


#     MAXIMUM TEMPERATURE AND HR PCN

HRMAXTEMPEGGPC <- slidingwin(xvar = list(Temp = PCENV$Max_Temp),
                            exclude = NA,
                            cdate = PCENV$NEWDATE,
                            bdate = PCEGG12$NEWDATE,
                            baseline = lmer(log10(HR) ~ YEAR + (1|NEST_BOX), data = PCEGG12),
                            cinterval = "day",
                            range = c(20, 0),
                            type = "relative",
                            stat = c("mean"),
                            func = c("lin","quad"))

HRMAXTEMPEGGPC$combos
summary(HRMAXTEMPEGGPC[[1]]$BestModel)

lm1=lm(yvar~factor(YEAR)+climate, data=HRMAXTEMPEGGPC[[1]]$BestModelData)
lmer1=lmer(yvar~factor(YEAR)+climate+(1|NEST_BOX), data=HRMAXTEMPEGGPC[[1]]$BestModelData)
anova(lmer1,lm1)

plotwin(dataset=HRMAXTEMPEGGPC[[1]]$Dataset)
plot(HRMAXTEMPEGGPC[[1]]$BestModelData)

Egg2 <- HRMAXTEMPEGGPC[[1]]$Dataset
summary(Egg2)

RandEgg <- randwin(repeats = 1000, 
                   window = "sliding", 
                   xvar=list(Temp = PCENV$Max_Temp), 
                   cdate=PCENV$NEWDATE,
                   bdate = PCEGG12$NEWDATE,
                   baseline= lmer(log10(HR) ~ YEAR + (1|NEST_BOX), data = PCEGG12), 
                   stat= "mean", 
                   range=c(20,0), 
                   func="lin",
                   type ="relative", 
                   cinterval = "day")

plotall(dataset = HRMAXTEMPEGGPC[[1]]$Dataset, datasetrand = RandEgg[[1]],
        cw1 = 0.95, cw2 = 0.5, cw3 = 0.25,
        title = NULL, arrow = FALSE)

res_sim <- DHARMa::simulateResiduals(HRMAXTEMPEGGPC[[1]]$BestModel, re.form = ~ (1|NEST_BOX))
plot(res_sim)

#     MINIMUM TEMPERATURE AND HR PCN ###############

HRMINTEMPEGGPC <- slidingwin(xvar = list(Temp = PCENV$Min_Temp),
                             exclude = NA,
                             cdate = PCENV$NEWDATE,
                             bdate = PCEGG12$NEWDATE,
                             baseline = lmer(log10(HR) ~ YEAR+(1|NEST_BOX), data = PCEGG12),
                             cinterval = "day",
                             range = c(20, 0),
                             type = "relative",
                             stat = c("mean"),
                             func = c("lin","quad"))

HRMINTEMPEGGPC$combos
summary(HRMINTEMPEGGPC[[1]]$BestModel)

lm1=lm(yvar~factor(YEAR)+climate+I(climate^2), data=HRMINTEMPEGGPC[[1]]$BestModelData)
lmer1=lmer(yvar~factor(YEAR)+climate+I(climate^2)+(1|NEST_BOX), data=HRMINTEMPEGGPC[[1]]$BestModelData)
anova(lmer1,lm1)

plotwin(dataset=HRMINTEMPEGGPC[[1]]$Dataset)
plot(HRMINTEMPEGGPC[[1]]$BestModelData)

Egg2 <- HRMINTEMPEGGPC[[1]]$Dataset
summary(Egg2)

RandEgg <- randwin(repeats = 1000, 
                   window = "sliding", 
                   xvar=list(Temp = PCENV$Min_Temp), 
                   cdate=PCENV$NEWDATE,
                   bdate = PCEGG12$NEWDATE,
                   baseline= lmer(log10(HR) ~ YEAR + (1|NEST_BOX), data = PCEGG12),
                   stat= "mean", 
                   range=c(20,0), 
                   func="lin",
                   type ="relative", 
                   cinterval = "day")

plotall(dataset = HRMINTEMPEGGPC[[1]]$Dataset, datasetrand = RandEgg[[1]],
        cw1 = 0.95, cw2 = 0.5, cw3 = 0.25,
        title = NULL, arrow = FALSE)

res_sim <- DHARMa::simulateResiduals(HRMINTEMPEGGPC[[1]]$BestModel, re.form = ~ (1|NEST_BOX))
plot(res_sim)

PCEGG12$MIN_TEMP=HRMINTEMPEGGPC[[1]]$BestModelData$climate


#     RAIN AND HR PCN ###############

HRRAINPC <- slidingwin(xvar = list(Rain = PCENV$Acc_mill),
                             exclude = NA,
                             cdate = PCENV$NEWDATE,
                             bdate = PCEGG12$NEWDATE,
                             baseline = lmer(log10(HR) ~ YEAR+(1|NEST_BOX), data = PCEGG12),
                             cinterval = "day",
                             range = c(20, 0),
                             type = "relative",
                             stat = c("mean"),
                             func = c("lin","quad"))

HRRAINPC$combos
summary(HRRAINPC[[1]]$BestModel)

lm1=lm(yvar~factor(YEAR)+climate+I(climate^2), data=HRRAINPC[[1]]$BestModelData)
lmer1=lmer(yvar~factor(YEAR)+climate+I(climate^2)+(1|NEST_BOX), data=HRRAINPC[[1]]$BestModelData)
anova(lmer1,lm1)

plotwin(dataset=HRRAINPC[[1]]$Dataset)
plot(HRRAINPC[[1]]$BestModelData)

Egg2 <- HRRAINPC[[1]]$Dataset
summary(Egg2)

RandEgg <- randwin(repeats = 1000, 
                   window = "sliding", 
                   xvar=list(Rain = PCENV$Acc_mill), 
                   cdate=PCENV$NEWDATE,
                   bdate = PCEGG12$NEWDATE,
                   baseline= lmer(log10(HR) ~ YEAR+(1|NEST_BOX), data = PCEGG12), 
                   stat= "mean", 
                   range=c(20,0), 
                   func="lin",
                   type ="relative", 
                   cinterval = "day")

plotall(dataset = HRRAINPC[[1]]$Dataset, datasetrand = RandEgg[[1]],
        cw1 = 0.95, cw2 = 0.5, cw3 = 0.25,
        title = NULL, arrow = FALSE)

res_sim <- DHARMa::simulateResiduals(HRRAINPC[[1]]$BestModel, re.form = ~ (1|NEST_BOX))
plot(res_sim) 

PCEGG12$RAIN=HRRAINPC[[1]]$BestModelData$climate

#     WIND AND HR PCN ###############

HRWINDPC <- slidingwin(xvar = list(Wind = PCENV$Av_ws_12),
                       exclude = NA,
                       cdate = PCENV$NEWDATE,
                       bdate = PCEGG12$NEWDATE,
                       baseline = lmer(log10(HR) ~ YEAR + (1|NEST_BOX), data = PCEGG12),
                       cinterval = "day",
                       range = c(20, 0),
                       type = "relative",
                       stat = c("mean"),
                       func = c("lin","quad"))

HRWINDPC$combos
summary(HRWINDPC[[2]]$BestModel)

lm1=lm(yvar~factor(YEAR)+climate+I(climate^2), data=HRWINDPC[[2]]$BestModelData)
lmer1=lmer(yvar~factor(YEAR)+climate+I(climate^2)+(1|NEST_BOX), data=HRWINDPC[[2]]$BestModelData)
anova(lmer1,lm1)

plotwin(dataset=HRWINDPC[[2]]$Dataset)
plot(HRWINDPC[[2]]$BestModelData)

Egg2 <- HRWINDPC[[2]]$Dataset
summary(Egg2)

RandEgg <- randwin(repeats = 1000, 
                   window = "sliding", 
                   xvar=list(Wind = PCENV$Av_ws_12), 
                   cdate=PCENV$NEWDATE,
                   bdate = PCEGG12$NEWDATE,
                   baseline= lmer(log10(HR) ~ YEAR+(1|NEST_BOX), data = PCEGG12),
                   stat= "mean", 
                   range=c(20,0), 
                   func="quad",
                   type ="relative", 
                   cinterval = "day")

plotall(dataset = HRWINDPC[[2]]$Dataset, datasetrand = RandEgg[[1]],
        cw1 = 0.95, cw2 = 0.5, cw3 = 0.25,
        title = NULL, arrow = FALSE)

res_sim <- DHARMa::simulateResiduals(HRWINDPC[[2]]$BestModel, re.form = ~ (1|NEST_BOX))
plot(res_sim)


PCEGG12$WIND=HRWINDPC[[2]]$BestModelData$climate
PCEGG12$WIND2=HRWINDPC[[2]]$BestModelData$climate^2

write.csv(PCEGG12, "EGG_DATA_CC12_PCN.csv", row.names = FALSE)


###   The below looks at both locations together but upon review it is better to analyse separately


############################################################
############################################################
############################################################

############      EGG HEART RATES AND CLIMATE BOTH LOCATIONS   ###########

############################################################
############################################################

setwd("X:\\01-BackupHD_230626\\Jan\\Joe Churchill\\chapter 3")

ENV=read.csv("ENV_ALL.csv")
ENV$NEWDATE <- as.Date(ENV$DATE,
                         format = "%d/%m/%Y")

EGG12_1=read.csv("EGG12_ALL.csv")
EGG12=subset(EGG12_1,HR!=0 & !is.na(HR))
EGG12$NEWDATE <- as.Date(EGG12$MEASURE_DATE,
                           format = "%d/%m/%Y")

hist(EGG12$HR)
EGG12
shapiro.test(EGG12$HR)


EGG12$NEST_BOX=paste(EGG12$NEST_BOX,EGG12$LOCATION,sep="")

####
#             AVERAGE TEMPERATURE AND HR ALL DATA
####

HRAVTEMPALL <- slidingwin(xvar = list(Temp = ENV$Av_temp),
                              exclude = NA,
                              cdate = ENV$NEWDATE,
                              bdate = EGG12$NEWDATE,
                              baseline = lmer(HR ~ YEAR + LOCATION +(1|NEST_BOX), data = EGG12),
                              cinterval = "day",
                              range = c(20, 0),
                              type = "relative",
                              stat = c("mean"),
                              func = c("lin","quad"), spatial = list(EGG12$LOCATION, ENV$LOCATION))

HRAVTEMPALL$combos
summary(HRAVTEMPALL[[1]]$BestModel)

plotwin(dataset=HRAVTEMPALL[[1]]$Dataset)
plot(HRAVTEMPALL[[1]]$BestModelData)

Egg2 <- HRAVTEMPALL[[1]]$Dataset
summary(Egg2)

RandEgg <- randwin(repeats = 1000, 
                   window = "sliding", 
                   xvar=list(Temp = ENV$Av_temp), 
                   cdate=ENV$NEWDATE,
                   bdate = EGG12$NEWDATE,
                   baseline= lmer(HR ~ YEAR + LOCATION +(1|NEST_BOX), data = EGG12),
                   stat= "mean", 
                   range=c(20,0), 
                   func="lin",
                   type ="relative", 
                   cinterval = "day", 
                   spatial = list(EGG12$LOCATION, ENV$LOCATION))

plotall(dataset = HRAVTEMPALL[[1]]$Dataset, datasetrand = RandEgg[[1]],
        cw1 = 0.95, cw2 = 0.5, cw3 = 0.25,
        title = NULL, arrow = FALSE)

climwin::plotdelta(HRAVTEMPALL[[1]]$Dataset)

####
#             MAXIMUM TEMPERATURE AND HR ALL DATA
####

HRMAXTEMPALL <- slidingwin(xvar = list(Temp = ENV$Max_Temp),
                               exclude = NA,
                               cdate = ENV$NEWDATE,
                               bdate = EGG12$NEWDATE,
                               baseline = lmer(HR ~ YEAR + LOCATION +(1|NEST_BOX), data = EGG12),
                               cinterval = "day",
                               range = c(20, 0),
                               type = "relative",
                               stat = c("max"),
                               func = c("lin","quad"), spatial = list(EGG12$LOCATION, ENV$LOCATION))

HRMAXTEMPALL$combos
summary(HRMAXTEMPALL[[1]]$BestModel)

plotwin(dataset=HRMAXTEMPALL[[1]]$Dataset)
plot(HRMAXTEMPALL[[1]]$BestModelData)

Egg2 <- HRMAXTEMPALL[[1]]$Dataset
summary(Egg2)

RandEgg <- randwin(repeats = 1000, 
                   window = "sliding", 
                   xvar=list(Temp = ENV$Max_Temp), 
                   cdate=ENV$NEWDATE,
                   bdate = EGG12$NEWDATE,
                   baseline= lmer(HR ~ YEAR + LOCATION +(1|NEST_BOX), data = EGG12),
                   stat= "max", 
                   range=c(20,0), 
                   func="lin",
                   type ="relative", 
                   cinterval = "day", 
                   spatial = list(EGG12$LOCATION, ENV$LOCATION))

plotall(dataset = HRMAXTEMPALL[[1]]$Dataset, datasetrand = RandEgg[[1]],
        cw1 = 0.95, cw2 = 0.5, cw3 = 0.25,
        title = NULL, arrow = FALSE)


####
#             MINIMUM TEMPERATURE AND HR ALL DATA
####

HRMINTEMPALL <- slidingwin(xvar = list(Temp = ENV$Min_Temp),
                               exclude = NA,
                               cdate = ENV$NEWDATE,
                               bdate = EGG12$NEWDATE,
                               baseline = lmer(HR ~ YEAR + LOCATION +(1|NEST_BOX), data = EGG12),
                               cinterval = "day",
                               range = c(20, 0),
                               type = "relative",
                               stat = c("min"),
                               func = c("lin","quad"), spatial = list(EGG12$LOCATION, ENV$LOCATION))

HRMINTEMPALL$combos
summary(HRMINTEMPALL[[2]]$BestModel)

plotwin(dataset=HRMINTEMPALL[[2]]$Dataset)
plot(HRMINTEMPALL[[2]]$BestModelData)

Egg2 <- HRMINTEMPALL[[2]]$Dataset
summary(Egg2)

RandEgg <- randwin(repeats = 1000, 
                   window = "sliding", 
                   xvar=list(Temp = ENV$Min_Temp), 
                   cdate=ENV$NEWDATE,
                   bdate = EGG12$NEWDATE,
                   baseline= lmer(HR ~ YEAR + LOCATION +(1|NEST_BOX), data = EGG12),
                   stat= "min", 
                   range=c(20,0), 
                   func="quad",
                   type ="relative", 
                   cinterval = "day", 
                   spatial = list(EGG12$LOCATION, ENV$LOCATION))

plotall(dataset = HRMINTEMPALL[[2]]$Dataset, datasetrand = RandEgg[[1]],
        cw1 = 0.95, cw2 = 0.5, cw3 = 0.25,
        title = NULL, arrow = FALSE)


##############################################################

##            HR AND RAINFALL ALL DATA

##############################################################


HRRAINALL <- slidingwin(xvar = list(Rain = ENV$Acc_mill),
                         exclude = NA,
                         cdate = ENV$NEWDATE,
                         bdate = EGG12$NEWDATE,
                         baseline = lmer(HR ~ YEAR + LOCATION +(1|NEST_BOX), data = EGG12),
                         cinterval = "day",
                         range = c(20, 0),
                         type = "relative",
                         stat = c("mean"),
                         func = c("lin","quad"), spatial = list(EGG12$LOCATION, ENV$LOCATION))

HRRAINALL$combos
summary(HRRAINALL[[1]]$BestModel)

plotwin(dataset=HRRAINALL[[1]]$Dataset)
plot(HRRAINALL[[1]]$BestModelData)

Egg3 <- HRRAINALL[[2]]$Dataset
summary(Egg3)

RandEgg1 <- randwin(repeats = 1000, 
                    window = "sliding", 
                    xvar=list(Rain = ENV$Acc_mill), 
                    cdate=ENV$NEWDATE,
                    bdate = EGG12$NEWDATE,
                    baseline= lmer(HR ~ YEAR + LOCATION +(1|NEST_BOX), data = EGG12),
                    stat= "mean", 
                    range=c(20,0), 
                    func="lin",
                    type ="relative", 
                    cinterval = "day", 
                    spatial = list(EGG12$LOCATION, ENV$LOCATION))

plotall(dataset = HRRAINALL[[1]]$Dataset, datasetrand = RandEgg1[[1]],
        cw1 = 0.95, cw2 = 0.5, cw3 = 0.25,
        title = NULL, arrow = FALSE)


################################################################

########            HR AND WIND SPEED ALL DATA

################################################################

HRWINDALL <- slidingwin(xvar = list(Wind = ENV$Av_ws_12),
                         exclude = NA,
                         cdate = ENV$NEWDATE,
                         bdate = EGG12$NEWDATE,
                         baseline = lmer(HR ~ YEAR + LOCATION +(1|NEST_BOX), data = EGG12),
                         cinterval = "day",
                         range = c(20, 0),
                         type = "relative",
                         stat = c("mean"),
                         func = c("lin","quad"), spatial = list(EGG12$LOCATION, ENV$LOCATION))

HRWINDALL$combos
summary(HRWINDALL[[2]]$BestModel)

plotwin(dataset=HRWINDALL[[2]]$Dataset)
plot(HRWINDALL[[2]]$BestModelData)

Egg4 <- HRWINDALL[[2]]$Dataset
summary(Egg4)

RandEgg2 <- randwin(repeats = 1000, 
                    window = "sliding", 
                    xvar=list(Wind = ENV$Av_ws_12), 
                    cdate=ENV$NEWDATE,
                    bdate = EGG12$NEWDATE,
                    baseline= lmer(HR ~ YEAR + LOCATION +(1|NEST_BOX), data = EGG12),
                    stat= "mean", 
                    range=c(20,0),
                    func="quad",
                    type ="relative", 
                    cinterval = "day", 
                    spatial = list(EGG12$LOCATION, ENV$LOCATION))

plotall(dataset = HRWINDALL[[2]]$Dataset, datasetrand = RandEgg2[[1]],
        cw1 = 0.95, cw2 = 0.5, cw3 = 0.25,
        title = NULL, arrow = FALSE)
```
