## Supplementary material for "Climatic conditions predict embryonic development in thorn tailed Rayadito (*Aphrastura spinicauda*)": Data and R scripts: EmbryonicDev_Chapter2_DataRepository.txt

author name: Joseph Eric Badji-Churchill
p-number: P310224
title of the thesis: The affect of climate change on Thorn-Tailed Rayadito's in Patagonia
year of publication: 2026
research group: Behavioural Physiology
promotor(s)/co-promotor(s): Prof. Jan Komdeur & Prof. Rodrigo Vasquez
contact person: Joseph Badji-Churchill [] or []

contents of the archive:
<list the chapters in the thesis and indicate whether a chapter has been published>

Chapter 2 - "Climatic conditions predict embryonic development in thorn tailed Rayadito (Aphrastura spinicauda)"
Chapter 3 - "Adaptation to climate driven environments in a Patagonian suboscine passerine"
Chapter 4 (published) - "The delayed effect of climatic conditions on pre-fledging nestling haematocrit in a suboscine species in Patagonia"

<list all files/folders and explain per folder/file what it contains>

Chapter 2 data:

ENV_DATA_ALL_1820.csv file contains all weather data in both locations (Navarino and Pucón) / input for R-script "Embryonic_Data_Modelling.R" and "Embryonic_Dev_D12Clim.R" / used to produce figures 2-5 and tables 1 and 2

ENV_DATA_NVRN_ALL.csv file contains all weather data in Navarino; input for R-script "Embryonic_Data_Modelling.R" and "Embryonic_Dev_D12Clim.R", used to produce figures 2-5 and tables 1 and 2

ENV_DATA_PCN_ALL.csv file contains all weather data in Pucón; input for R-script "Embryonic_Data_Modelling.R" and "Embryonic_Dev_D12Clim.R", used to produce figures 2-5 and tables 1 and 2

EGG_DATA_CC12.csv file contains all embryonic heart rate readings on the 12th day of incubation in both locations (Navarino and Pucón); input for R-script "Embryonic_Data_Modelling.R", used for statistical tests on between location comparisons

ALL_EGG_DATA.csv file contains all embryonic heart rate readings during incubation and egg mass/size measurements in both locations (Navarino and Pucón); input for R-script "Embryonic_Data_Modelling.R", used for statistical tests on between location comparisons and egg mass comparisons

EGG_DATA_CC12_NVRN.csv contains all embryonic heart rate readings on the 12th day of incubation in Navarino, used to produce figure 4 and tables 1 and 3; input for R-script "Embryonic_Data_Modelling.R" and "Embryonic_Dev_D12Clim.R", used for climwin analysis and LMERs in Navarino

EGG_DATA_CC12_PCN.csv contains all embryonic heart rate readings on the 12th day of incubation in Pucón, used to produce figures 2, 3 and 5 and tables 1 and 2; input for R-script "Embryonic_Data_Modelling.R" and "Embryonic_Dev_D12Clim.R", used for climwin analysis and LMERs in Pucón

Chapter 3 data:

OVERALL_GROWTH_NVRN.csv file containing growth rate and pre-fledging weights of nestlings in Navarino; input for R-script "FinalStatsNestlingGrowth.R" and "New_Climwin_Growth_Rates.R", used to produce figure 6 and tables 1 and 2 in Navarino

OVERALL_GROWTH_PCN.csv file containing growth rate and pre-fledging weights of nestlings in Pucón; input for R-script "FinalStatsNestlingGrowth.R" and "New_Climwin_Growth_Rates.R", used to produce tables 1 and 2 in Pucón

NESTLING_GROWTH_PER_DAY.csv file containing growth rate and pre-fledging weights of nestlings in Navarino and Pucón; input for R-script "FinalStatsNestlingGrowth.R" and "New_Climwin_Growth_Rates".R, used for analysis of comparisons between locations of growth rates

D1NVRN.csv file containing hatch weights of nestlings in Navarino; input for R-script "FinalStatsNestlingGrowth.R" and "New_Climwin_Growth_Rates.R", used to produce figures 4 and 5 and tables 1 and 2 in Navarino

D1PCN.csv file containing hatch weights of nestlings in Pucón; input for R-script "FinalStatsNestlingGrowth.R" and "New_Climwin_Growth_Rates.R", used to produce figures 2 and 3 and tables 1 and 2 in Navarino

D1ALL.csv file containing hatch weights of nestlings in Navarino and Pucón; input for R-script "FinalStatsNestlingGrowth.R" and "New_Climwin_Growth_Rates.R", used for analysis of comparisons between locations of hatching weights

Chapter 4:

HD1.csv file containing all physiological measurements and haematocrit measurements for nestlings on Day 16; input for R-script "Haematocrit code", used to produce figures 2A, 2B, 3, 4A and 4B and table 1.

ENV_DATA_ALL_1820.csv file containing weather data for both locations and in both years; input for R-script "Haematocrit code", used for climwin analysis analysing locations together

TEMP_18_20_NVPC.csv file containing temperature data for both locations in both years; input for R-script "Haematocrit code", used for climwin analysis and LMERs to compare between locations

1920_DATA.csv file containing isolated data from HD1 for 2019-2020 (1920) season only, this was used to analyse Habitat quality separately (see manuscript); input for R-script "Haematocrit code", used for graph making of temperature and rain


<document any relevant info about the archive, e.g. special conditions for data accessibility, generic databases or datasets that are used in more than one chapter (no need to archive these in each chapter, just once will do), any other relevant info>

None to document

_____________________________________________________________________________

EmbryonicDev_Chapter2_DataRepository.txt: "Climatic conditions predict embryonic development in thorn tailed Rayadito (Aphrastura spinicauda)"

Unpublished: "Climatic conditions predict embryonic development in thorn tailed Rayadito (Aphrastura spinicauda)"

<>
<>
<>

Contents of chapter 2: "Climatic conditions predict embryonic development in thorn tailed Rayadito (Aphrastura spinicauda)"

Data:
<list all data files/folders and explain per folder/file what it is, and if needed, how to use it>

ENV_DATA_ALL_1820.csv file contains all weather data in both locations (Navarino and Pucón) / input for all R-scripts used in chapter 2 and 3 (please refer to "code" section below) / Not listed in other metafiles as instructed

ENV_DATA_NVRN_ALL.csv file contains all weather data in Navarino / input for all R-scripts used in chapter 2 and 3 (please refer to "code" section below) / Not listed in other metafiles as instructed

ENV_DATA_PCN_ALL.csv file contains all weather data in Pucón / input for all R-scripts used in chapter 2 and 3 (please refer to "code" section below) / Not listed in other metafiles as instructed

EGG_DATA_CC12.csv file contains all embryonic heart rate readings on the 12th day of incubation in both locations (Navarino and Pucón) / input for R-script "Embryonic_Data_Modelling.R", used for statistical tests on between location comparisons

ALL_EGG_DATA.csv file contains all embryonic heart rate readings during incubation and egg mass/size measurements in both locations (Navarino and Pucón); input for R-script "Embryonic_Data_Modelling.R", used for statistical tests on between location comparisons and egg mass comparisons

EGG_DATA_CC12_NVRN.csv contains all embryonic heart rate readings on the 12th day of incubation in Navarino / input for R-script "Embryonic_Data_Modelling.R" and "Embryonic_Dev_D12Clim.R", used for climwin analysis and LMERs in Navarino

EGG_DATA_CC12_PCN.csv contains all embryonic heart rate readings on the 12th day of incubation in Pucón / input for R-script "Embryonic_Data_Modelling.R" and "Embryonic_Dev_D12Clim.R", used for climwin analysis and LMERs in Pucón

Code:
<list all code/scripts files/folder and explain per folder/file what it is, and if needed, how to use it>

Embryonic Dev D12Clim.R / The script containing the code for the "climwin" package for analysis of climate data and embryonic development / to load data, refer to Embryonic Data Modelling.R script

Embryonic Data Modelling.R / The script containing the code for further analysis of data using LMERs and other statistical tests, the data generated from Embryonic Dev D12Clim.R is used here 

Documents:
<list all document files/folders and explain per folder/file what it is, and if needed, how to use it>

No additional documents

Software:
<list all non-standard software that is used with program/package version and explain how to obtain it (link), and if needed, how to use it>

RStudio / R-4.5.2

External data:
<list all external datasets used and explain how to access them (link, contact details of a data manager, etc)>

Weather data in ENV_DATA_ALL_1820.csv, ENV_DATA_NVRN_ALL.csv and ENV_DATA_PCN_ALL.csv all extracted from weather stations "Pucon Aerorome, Pucon" and "Guardiamarina Zanatu Airport, Cpae Horn" [https://agrometeorologia.cl/]
